# Breastfeeding Enables Systemic Transfer of Functional Maternal Immune Cells to Neonates

**DOI:** 10.64898/2026.11.17.767284

**Authors:** Si Naftaly, Tamar Gross Lev, Almog Angel, Lilac Haimovich-Caspi, Shir Erez, Chen Yanovich-Ben-Uriel, Michal Brunwasser-Meirom, Dvir Aran, Assaf Zinger

## Abstract

Does breastfeeding transfer living maternal immune cells that persist and function in offspring, beyond providing soluble immune factors? Using a foster-nursing mouse model that eliminates prenatal exposure, this study directly tests whether maternal cells can be transferred postnatally through breastmilk. We found that intact maternal immune cells survive neonatal gastrointestinal transit, enter the circulation, and disseminate to peripheral organs. Single-cell RNA sequencing shows that transferred cells span multiple hematopoietic lineages, dominated by monocyte-derived macrophages with a distinct stress-adapted, inflammatory transcriptional state compatible with persistence in the neonatal environment. Interestingly, these maternal cells are detected at sites of neonatal tissue injury, consistent with chemotactic responsiveness. Our findings suggest that breastfeeding may serve as a postnatal route for maternal cellular transfer, potentially extending maternal immune influence beyond gestation.

## Introduction

Breastfeeding is a defining feature of early postnatal life and a major determinant of neonatal immune development. Its protective effects against infection and inflammation have traditionally been attributed to the transfer of soluble factors, including antibodies, cytokines, growth factors, and antimicrobial molecules^1,2^. These components shape immune maturation, barrier function, and host– microbe interactions during a critical developmental window^3–5^. However, whether breastfeeding also mediates the postnatal transfer of viable maternal immune cells that persist and function within the offspring has remained unclear.

The concept that breastfeeding conveys maternal immune protection dates back more than a century^6,7^. Breastmilk contains abundant viable maternal cells, including leukocytes, epithelial cells, and progenitor populations derived from both the mammary gland and systemic circulation^8–11^. Early experimental studies suggested that some of these cells may cross the neonatal gastrointestinal tract and localize to peripheral organs. For example, radiolabeled leukocyte experiments detected radioactive signals in the spleen, liver, and bone marrow of premature baboons after feeding^12^, while studies in lambs and piglets reported migration of viable milk-derived lymphocytes into the intestinal mucosa^13^. More recently, transgenic mouse models expressing green fluorescent protein (GFP) have enabled direct visualization of maternal cell transfer. Using this approach, GFP-expressing (GFP^⁺^) maternal T cells have been detected in the intestinal epithelium of nursing pups^14,15^, and GFP^⁺^ cells have been reported in the livers and brains of offspring nursed by GFP^⁺^ dams^16,17^. Consistent with these observations, Twigger et al. detected breastmilk-derived maternal cells in the brains of murine offspring and adopt neural-like phenotypes^18^. In addition, maternally derived immune cells transferred during breastfeeding have been shown to confer protection against parasitic infection in nursing offspring^19^.

Maternal–offspring cellular exchange is well established during pregnancy, where bidirectional trafficking of cells across the placenta leads to long-lived microchimerism in both mother and child. Maternal cells acquired in utero can persist for years and have been implicated in immune modulation, tissue repair, and disease susceptibility^20–25^. In contrast, postnatal cellular transfer has received far less attention^26,27^. Recent single-cell transcriptomic studies have further characterized the cellular diversity of human breastmilk, identifying distinct lactocyte and immune cell populations across lactation stages^10,28,29^. These advances, together with reviews highlighting the broader role of maternal leukocytes in infant immune programming^30,31^, have provided a detailed molecular catalog of milk-borne cells. Yet whether intact maternal immune cells enter neonatal circulation and acquire functional roles within the offspring remains unresolved.

A central challenge in defining postnatal maternal cell transfer has been distinguishing viable cellular engraftment from passive uptake of cell fragments, apoptotic bodies, or extracellular vesicles. Indeed, Doerfler et al. recently demonstrated that maternal milk cell components can be taken up by infant liver macrophages via extracellular vesicle-mediated transport, highlighting the complexity of distinguishing intact cell transfer from cell-free material exchange^32^. As a result, it remains unknown whether breastfeeding contributes to neonatal immunity through an active postnatal cellular pathway distinct from gestational cell exchange^23,33^.

Addressing this question requires unambiguous tracking of maternal cells after birth and molecular characterization of their identity and functional state within the neonatal host. In this study, we asked whether breastfeeding provides a postnatal route for the transfer of intact maternal immune cells into the offspring and whether such cells remain biologically responsive within the neonatal environment. To address this, we developed a foster-nursing mouse model that excludes prenatal exposure and enables unambiguous tracking of maternally derived cells after birth. By integrating *in vivo* cell tracing with flow cytometry, tissue imaging, and single-cell transcriptomic profiling, we examined the fate, identity, and molecular state of maternal cells following gastrointestinal passage and systemic entry into the neonate. This approach allowed us to determine whether breastmilk-derived maternal immune cells persist beyond the gut, how they differ from endogenous neonatal immune populations, and whether they retain the capacity to respond to inflammatory cues within neonatal tissues.

Our findings provide evidence that viable maternal immune cells transferred through breastfeeding are detected systemically and at sites of neonatal tissue injury, with stress-adapted maternal macrophages representing the dominant transferred population, supporting the concept that breastfeeding may represent an active cellular pathway contributing to the neonatal immune landscape.

The pathophysiological relevance of postnatal maternal cell transfer remains poorly understood, yet the neonatal period represents one of the most immunologically vulnerable windows in human life. Neonates are born with an immature immune system that must rapidly adapt to microbial colonization, barrier maturation, and the transition from sterile in utero conditions to the antigen-rich extrauterine environment. Epidemiological evidence consistently links breastfeeding to reduced incidence and severity of neonatal sepsis^34^, necrotising enterocolitis^35^, respiratory infections^36^, and inflammatory bowel disease^37^ during infancy and benefits that extend beyond what passive antibody transfer alone can explain. If intact maternal immune cells traverse the neonatal gut, enter circulation, and home to sites of inflammation, they could constitute an active cellular component of this protection: providing innate immune reinforcement at a time when neonatal myeloid responses are functionally limited, contributing myeloid effectors capable of sensing pathogen-associated cues and coordinating early inflammatory resolution, or shaping the neonatal immune landscape through direct cell-cell interactions with tissue-resident and circulating immune populations. In addition, dysregulation of this pathway, for example under conditions of maternal infection, inflammation, or altered lactation could theoretically contribute to neonatal immune imbalance, allergy predisposition, or inflammatory pathology. Understanding whether and how breastmilk-derived maternal cells participate in neonatal immunity therefore has direct implications for interpreting the epidemiology of breastfeeding-associated health outcomes and for identifying novel cellular mechanisms that shape neonatal immune development.

## Results

### A foster-nursing model enables unambiguous tracking of postnatal maternal cell transfer

Breastfeeding has been proposed as a route for the postnatal transfer of viable maternal cells to the offspring; however, whether such cells survive gastrointestinal passage, access neonatal tissues, and persist systemically remain undefined. To determine whether breastfeeding enables postnatal transfer of intact, viable maternal cells into the neonatal host, we employed a foster-nursing approach in which GFP^+^ transgenic dams nursed wild-type (WT; GFP^-^) pups immediately after delivery. GFP^+^ mice ubiquitously express GFP under the human ubiquitin C promoter, enabling direct visualization and tracking of maternally derived cells within the offspring^38^.

WT neonates were fostered immediately after birth and maintained with GFP^⁺^ dams for 18 days, ensuring continuous exposure to breastmilk-derived maternal cells. GFP^⁺^ maternal cells were subsequently analyzed by flow cytometry and GFP+ cells detected in the circulation further characterized by single-cell RNA sequencing (scRNA-seq) to define their cellular identities and potential functional states (**Fig. 1A**). This GFP^⁺^/GFP^⁻^ lineage-tracing strategy enabled direct discrimination of maternally derived cells from offspring-derived cells, as any GFP^⁺^ cells detected in recipient tissues could only originate from the foster dam following postnatal transfer through lactation.

**Figure 1.**
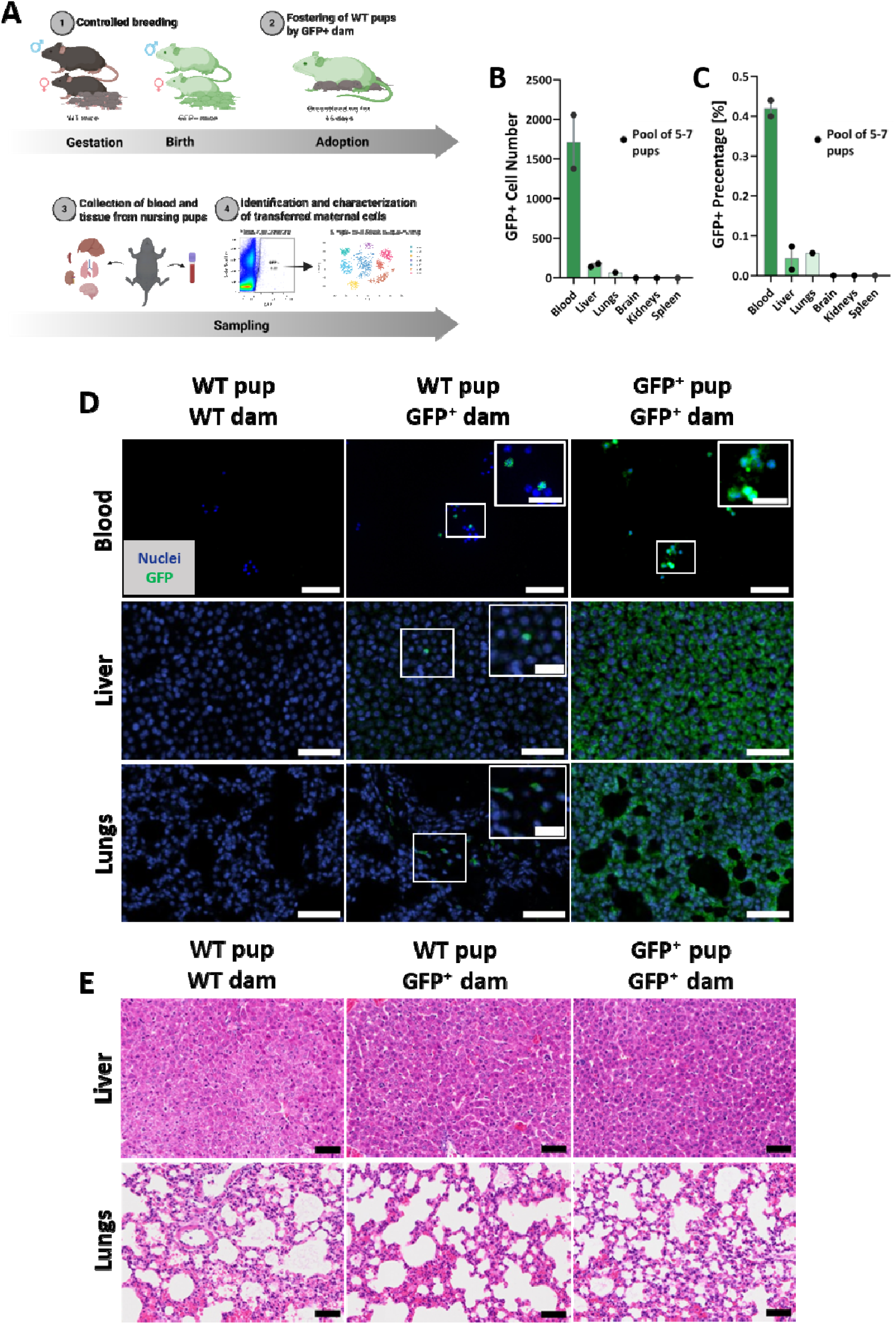
Breastmilk maternal cells are detected in nursing offspring circulation and peripheral organs. **(A)** Schematic overview of the foster-nursing strategy used to track maternal cell transfer. WT pups were nursed by GFP^+^ dams to enable identification of breastmilk-derived maternal cells. Flow cytometric quantification showing the **(B)** frequency and **(C)** absolute number of GFP^+^ maternal cells in peripheral blood and indicated organs of WT pups nursed by GFP^+^ dams. Representative images from WT pups nursed by WT dams (negative control), WT pups nursed by GFP^+^ dams, and GFP^+^ pups nursed by GFP^+^ dams (positive control) of **(D)** blood, liver and lungs stained for nuclei (blue) and GFP (green) or **(E)** liver and lungs stained for H&E, imaged using an Olympus Slideview VS200. Scale bars: 50 µm (overview images); 20 µm (magnified images). Each dot represents one independently processed pooled sample of five to seven pups from a single litter; n = 1-2 independent litters. Results are shown as mean ± SEM.

### Maternal cells circulate in nursing pups

Using flow cytometry, we quantified the presence and distribution of GFP^⁺^ maternal cells in the blood and selected organs of 18-day nursed WT pups from two independent litters. Peripheral blood was collected, subjected to red blood cell (RBC) lysis, and processed for blood cells isolation. The liver, lungs, brain, kidneys, and spleen were mechanically dissociated, followed by RBC lysis of the resulting cell suspensions. After exclusion of debris, doublets, and non-viable events using Zombie Violet staining and a stringent gating strategy (**Fig. S1A**), GFP^⁺^ maternal cells were readily detected in the circulation, with lower but measurable frequencies in the liver and lungs. Back-gating demonstrates that GFP^⁺^ events localize within the expected live-cell population and were not distributed among debris or excluded events (**Fig. S1B**). In contrast, GFP^⁺^ cells were below the limit of detection in the brain, kidney, or spleen (**Fig. 1B**). Quantitatively, approximately 1.7×10³±3×10² GFP^⁺^ cells were identified in circulating blood, corresponding to 0.42±0.02% of blood cells, compared with 0.04±0.03% and 0.06% in liver and lungs, respectively (**Fig. 1B-C**). These findings provide direct evidence that maternal cells delivered postnatally through breastfeeding gain access to the neonatal bloodstream and disseminate into select peripheral organs, establishing a postnatal pathway of maternal cell transfer independent of gestation.

To further validate our flow cytometry findings and determine the spatial localization of maternal cells within intact tissues, we stained blood cells, liver and lung histological sections with an anti-GFP antibody and the nuclear marker DAPI. Given the low abundance of transferred maternal cells in recipient tissues, GFP-positive cells were detected by anti-GFP immunostaining to maximize detection sensitivity. Consistent with our flow cytometry results, antibody-based GFP labeling revealed GFP^⁺^ cells in blood and organs of WT pups nursed by GFP^⁺^ dams, while no GFP signal was detected in WT pups nursed by WT dams (negative control), confirming that the observed GFP^⁺^ cells are not artifacts of cytometric detection. This assay was further validated by strong GFP staining in GFP^+^ pups nursed by GFP^+^ dams (positive control; **Fig. 1D**). Tissue morphology and overall architectural integrity were comparable across all groups (**Fig. 1E**). The enrichment of GFP^⁺^ cells in pup blood, coupled with their lower but reproducible presence in peripheral organs, demonstrates that breastmilk-derived maternal cells disseminate through the neonatal circulation and access multiple organs, raising the possibility of their persistence or integration within neonatal tissues.

### Maternal cells localize to the neonatal intestinal mucosa

Previous studies have reported that breastmilk-derived maternal immune cells can be detected within the intestine of nursing offspring, particularly in the intestinal epithelium and Peyer’s patches^14,15^. To determine whether maternal cells are similarly localized within the neonatal intestine in our foster-nursing model, we examined intestinal sections from WT pups nursed by GFP^⁺^ dams. H&E staining first confirmed the preserved intestinal architecture, with well-defined villi and epithelial structures across the examined tissue sections (**Fig. 2A**). Fluorescence imaging revealed GFP^⁺^ maternal cells within the intestinal mucosa, with GFP signal detected in close association with the epithelial compartment, whereas no GFP signal was observed in WT pups nursed by WT dams (**Fig. 2B**). The corresponding individual fluorescence channels further resolved the distribution of nuclei, E-cadherin, CD31, and GFP, confirming the localization of GFP^⁺^ cells within the mucosal tissue (**Fig. 2B**).

**Figure 2.**
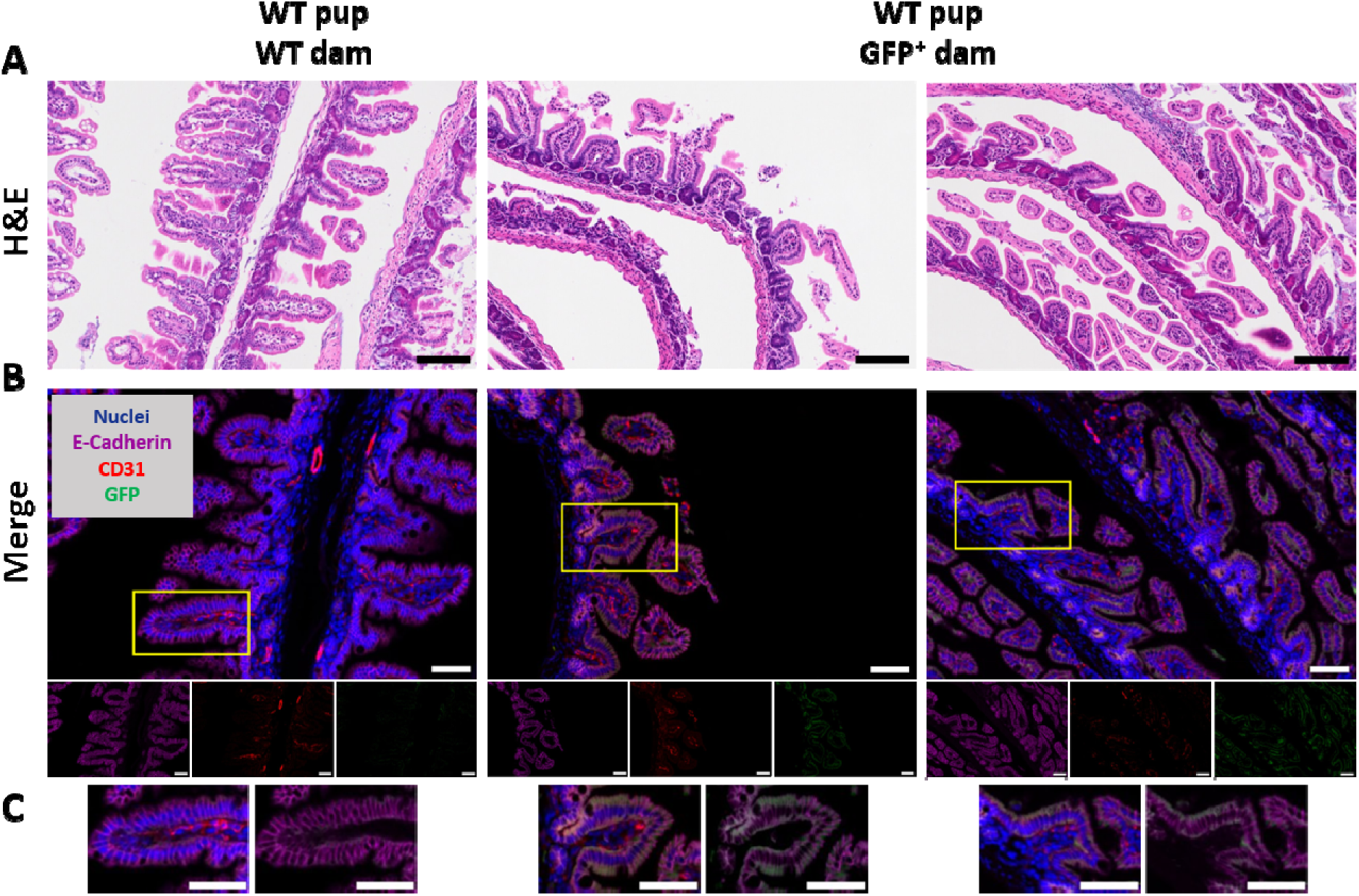
Maternal cells transferred through breastfeeding localize within the intestinal mucosa of nursing offspring. Representative histological and immunofluorescence images of intestinal tissue from WT pups nursed by GFP^⁺^ dams, demonstrating the presence and localization of breastmilk-derived maternal cells. **(A)** Representative H&E-stained sections showing the intestinal architecture and mucosal organization. Scale bars: 100 µm. **(B)** Representative merged immunofluorescence images showing nuclei (blue), E-cadherin (magenta) marking intestinal epithelial cells, CD31 (red) marking endothelial cells, and GFP (green) identifying maternal cells transferred through breast milk. Yellow boxes indicate regions shown at higher magnification. Scale bars: 50 µm. **(C)** Higher-magnification images of the indicated regions demonstrating GFP^⁺^ maternal cells in proximity to the intestinal epithelium and underlying vascular compartment. Scale bars: 50 µm.

Higher-magnification views of the boxed regions showed GFP^⁺^ signal closely associated with E-cadherin-positive epithelial structures and not obviously restricted to the intercellular spaces, whereas CD31-positive vasculature was localized beneath the epithelial layer (**Fig. 2C**). This spatial distribution is consistent with maternal cells being closely associated with the epithelial compartment, although the available two-dimensional images do not permit definitive discrimination between intracellular and paracellular localization. Together, these observations demonstrate that maternally derived cells are present within the neonatal intestinal mucosa and positioned near the epithelial– vascular interface, supporting the intestine as a site of maternal cell localization in nursing offspring.

### Circulating GFP^⁺^ cells are significantly enriched in pups nursed by GFP^⁺^ dams

To characterize the specific breastmilk-derived maternal cell populations transferred to neonates, we isolated GFP^+^ cells from neonatal blood using fluorescence-activated cell sorting (FACS). Blood from WT pups nursed by WT dams served as a stringent negative control for establishing the GFP-positive gate and defining background autofluorescence. Non-cellular events and debris were excluded using a standardized gating strategy (**Fig. S1C**). Across biological replicates, WT pups nursed by GFP^⁺^ dams exhibited a significant 2.2-fold enrichment of circulating GFP^⁺^ events compared with WT pups nursed by WT dams (0.29 ± 0.07% vs. 0.13 ± 0.03%, p=0.0323; **Fig. 3A**), confirming the maternal origin of these circulating cells. Representative flow cytometry plots are shown in Supplementary Fig. **S1C–D**. In the representative experiment, GFP^⁺^ events comprised 0.22% (804/358,888 events) of live, singlet cells in WT pups nursed by GFP^⁺^ dams, compared with 0.063% (203/322,337 events) in WT pups nursed by WT dams. Importantly, back-gating analysis demonstrated that GFP^⁺^ events localized within the expected cell population rather than debris or excluded events (**Fig. S1D**), supporting that these events represent genuine GFP^⁺^ cellular events rather than nonspecific background fluorescence or debris. These findings indicate that maternal GFP^⁺^ cells are transferred to the circulation of nursing pups. Sorted GFP^⁺^ cells were subsequently subjected to scRNA-seq to define their transcriptional identities and potential functional states.

**Figure 3.**
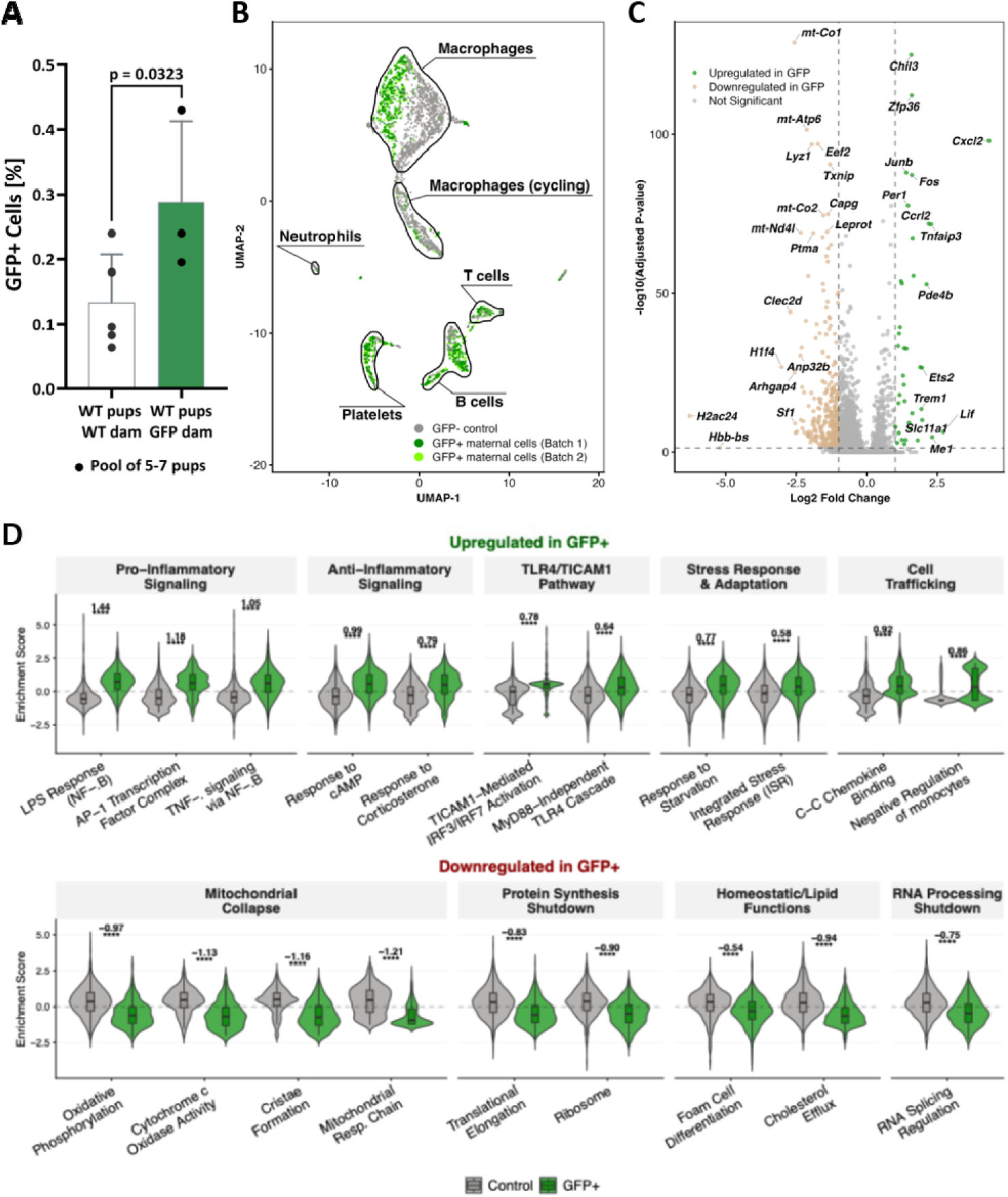
Single-cell transcriptomic profiling reveals a distinct molecular identity of transferred maternal cells. GFP^⁺^ cells from peripheral blood were purified by FACS. **(A)** Frequencies of circulating GFP^⁺^ cells in WT pups nursed by GFP^+^ dams compared with WT pups nursed by WT dams (negative control). Because the two groups consisted of independent animals, a one-tailed unpaired Student’s t-test was used; Data are presented as mean ± SEM; n = 3-5 independent litters per group. P value ≤ 0.05 among means was considered statistically significant. **(B–D)** Single-cell analysis was performed using two independent GFP^⁺^ biological replicates and one GFP^⁻^ control sample. **(B)** UMAP visualization of 2,818 cells colored by sample origin (gray = GFP^⁻^ control; light green = GFP^⁺^ maternal cells batch 1; dark green = GFP^⁺^ maternal cells batch 2). Black outlines denote cell-type clusters identified at resolution 0.1, with major populations labeled and corresponding cluster numbers indicated in parentheses (macrophages [C0], B cells [C1], cycling macrophages [C2], platelets [C3], T cells [C4], neutrophils [C5]). An additional minor cluster was present but not labeled for clarity. The macrophage cluster (C0) shows clear segregation between GFP^⁺^ maternal cells and GFP^⁻^ pup cells. **(C)** Volcano plot showing differential gene expression between GFP^⁺^ maternal macrophages and GFP^⁻^ control macrophages in cluster 0. Significance thresholds: adjusted p-value < 0.05, |log□FC| > 1. **(D)** Violin plots showing enrichment scores for key pathways differentially regulated between GFP^⁺^ maternal macrophages (green) and GFP^⁻^ control macrophages (gray). Upper panel: pathways upregulated in maternal cells. Lower panel: pathways downregulated in maternal cells. Numbers above violins indicate Cohen’s d effect size. Statistical significance: ****p < 0.0001 (Wilcoxon rank-sum test with FDR correction).

### Single-cell profiling reveals diverse hematopoietic lineages among transferred maternal cells

Using the 10x Genomics Chromium Single Cell Gene Expression Flex platform, we initially profiled GFP^⁺^ maternal cells from two independent biological replicates together with GFP^⁻^ control cells from age-matched pups nursed by WT dams. After quality control filtering and doublet removal, a total of 1,428 GFP^⁺^ maternal cells and 1,390 GFP^⁻^ control cells were retained for downstream analysis. By applying the standard Seurat analytic pipeline^39^ we identified seven transcriptionally distinct cell clusters (**Fig. 3B**). Cell type annotation based on SingleR^40^ and canonical markers revealed diverse hematopoietic lineages, including B cells (Cd79a^⁺^), T cells (Cd3d^⁺^), platelets (Pf4^⁺^), neutrophils (S100a9^⁺^), and multiple monocyte/macrophage populations ^40^ (**Fig. S2**). Cell type assignments were made from canonical markers within our own primary data; the published lactating mammary gland dataset was used only for the macrophage comparison presented in **Fig. S4** and did not contribute to these annotations. Platelet identity was supported by near-universal detection of Itga2b, Gp1ba and Pf4 within the corresponding cluster, together with Ppbp and Mpl. Because platelets derive from megakaryocytes and retain megakaryocytic transcripts, these markers do not define a separate megakaryocyte population, and no such cluster was identified. We note that platelet and megakaryocytic transcripts are particularly susceptible to ambient RNA, and none of the conclusions of this study depend on this cluster.

### Maternal macrophages are transcriptionally distinct

Notably, the largest transcriptional population (Cluster 0; **Fig. S2**) consisted of monocyte/macrophage-lineage cells, characterized by the expression of canonical myeloid markers such as Lyz2 and Adgre1 (**Fig. S2**). This cluster included 926 GFP^⁻^ and 610 GFP^⁺^ maternal cells, together accounting for 54.5% of all analyzed cells (**Fig. 3B**). Within this cluster, GFP^⁺^ maternal cells were transcriptionally distinct from GFP^⁻^ pup cells, indicating that transferred maternal macrophages adopt a transcriptional state that is clearly separable from endogenous pup cells of the same lineage. The additional macrophage cell cluster (Cluster 2; **Fig. 3B**) was predominantly composed of GFP^⁻^ pup cells, displaying proliferative markers such as Mki67 and Top2a, consistent with active cell cycling (**Fig. S2**). In contrast, GFP^⁺^ maternal cells were markedly underrepresented in this cycling population, suggesting limited proliferative capacity following transfer.

### Maternal macrophages show a distinct differential expression signature

We focused subsequent analysis on the primary macrophage cell cluster (Cluster 0) to elucidate the molecular characteristics distinguishing transferred maternal cells from endogenous pup macrophages. Differential expression analysis identified 39 significantly upregulated and 331 significantly downregulated genes in GFP^⁺^ cells compared to GFP^⁻^ control cells (adjusted p-value < 0.05, |log2 fold-change| > 1). Among the most highly upregulated genes in maternal cells were chemokine Cxcl2, atypical chemokine receptor Ccrl2, TNF-induced protein Tnfaip3, chitinase-like protein Chil3, transcription factor Fos, and zinc finger protein Zfp36 (**Fig. 3C**). Conversely, downregulated genes included mitochondrial-encoded transcripts (mt-Co1, mt-Atp6, mt-Nd4l), lysozyme Lyz1, and C-type lectin receptor Clec2d, among others.

### Inflammatory and stress programs dominate

Gene set enrichment analysis revealed that transferred maternal macrophages exhibit a distinct transcriptional profile characterized by inflammatory priming coupled with metabolic restriction and stress adaptation. To assess redundancy among enriched pathways, we constructed a gene-set similarity network based on shared genes across pathways. This analysis revealed that major functional categories form discrete clusters with limited gene overlap, indicating that the observed enrichments represent distinct transcriptional programs rather than repeated detection of the same genes (**Fig. S3**). The analysis showed that maternal cells displayed robust enrichment for pro-inflammatory signaling pathways, including LPS response signatures (Cohen’s d = 1.44, p < 0.0001), AP-1 transcription factor complex activation (d = 1.18, p < 0.0001), and TNFa signaling via NF-κB (d = 1.05, p < 0.0001) (**Fig. 3D**, upper panel). This inflammatory activation was accompanied by specific enrichment of the TICAM1-mediated TLR signaling pathway (d = 0.78, p < 0.0001) and MyD88-independent TLR4 cascade (d = 0.64, p < 0.0001), consistent with cells that may have encountered microbial components during gastrointestinal passage. Additionally, maternal cells showed enrichment for stress-adaptive responses, including responses to cAMP (d = 0.99, p < 0.0001) and corticosterone (d = 0.79, p < 0.0001), alongside signatures of starvation response (d = 0.77, p < 0.0001).

### Metabolic quiescence with immune readiness

In contrast to this inflammatory and stress-responsive phenotype, GFP^+^ maternal cells exhibited marked suppression of core metabolic and biosynthetic machinery (**Fig. 3D**, lower panel). We observed significant downregulation of protein synthesis pathways, including ribosomal function (d = -0.90, p < 0.0001) and translational elongation (d = -0.83, p < 0.0001). Mitochondrial function was similarly compromised, with reduced expression of electron transport chain components and suppression of mitochondrial cristae formation (d = -1.16, p < 0.0001). Lipid homeostasis pathways were also downregulated, including cholesterol efflux mechanisms (d = -0.94, p < 0.0001) and RNA splicing regulation (d = -0.75, p < 0.0001). This metabolic quiescence suggests that transferred cells prioritize survival and stress resistance overgrowth and proliferation.

### Maternal macrophages retain chemotaxis

Despite their metabolically restricted state, maternal cells retained functional capabilities for tissue navigation and immune coordination. We observed significant enrichment for C-C chemokine binding (d = 0.92, p < 0.0001) and negative regulation of monocytes (d = 0.86, p < 0.0001), consistent with maintained chemotactic responsiveness (**Fig. 3D**, upper panel). Among the most highly upregulated genes in maternal cells was Cxcl2, a chemokine associated with neutrophil recruitment and inflammatory coordination. Collectively, these findings indicate that transferred maternal cells adopt a stress-adapted phenotype: metabolically quiescent and stress-adapted yet immunologically primed to respond to inflammatory cues and navigate tissue environments.

This stress-adapted transcriptional profile suggests that circulating maternal macrophages occupy a distinct transcriptional state. To determine whether this state resembles the macrophage state described in the lactating mammary gland, we integrated our macrophage dataset with the published liMac macrophage dataset^41^ and compared gene-set scores across groups (**Fig. S4**). Control and GFP^⁺^ circulating macrophages showed comparable scores for a macrophage core gene set, confirming shared lineage identity with liMac clusters. However, circulating macrophages showed low scores for the liMac gene set defined in the original study, whereas a subset of liMac clusters exhibited high liMac gene-set scores. In parallel, GFP^⁺^ macrophages were enriched for the GFP^⁺^-upregulated signature derived from our dataset and showed reduced scores for the GFP^⁺^-downregulated signature, relative to control macrophages. liMac clusters did not mirror this combined pattern: they generally showed high scores for the GFP^⁺^-downregulated signature and limited enrichment of the GFP^⁺^-upregulated signature, with at most partial overlap in one subset. Thus, no liMac cluster recapitulated the paired high “up” and low “down” signature profile that defines circulating GFP^⁺^ macrophages, supporting the conclusion that circulating maternal macrophages represent a distinct macrophage state rather than a mobilized form of liMacs.

### Transgene detection, cell quality, and replicate concordance support the single-cell analysis

Because the single-cell probe panel included three custom probes targeting EGFP, transgene transcripts could be assessed directly from the sequencing data. EGFP transcripts were detected in 17 of 1,428 cells sorted from pups nursed by GFP^⁺^ dams, across both biological replicates, compared with none of 1,390 cells sorted from pups nursed by WT dams (Fisher’s exact test, p = 1.5 × 10^⁻^; **Fig. S5A, B**). EGFP detection was sparse, and EGFP^⁺^ cells could not therefore be assigned unequivocally to maternal origin based on transgene detection alone. Some EGFP^⁺^ cells also contained Y-chromosome transcripts.

The transferred macrophages exhibited a stress-associated transcriptional program, prompting us to assess whether this phenotype could instead reflect poor cell quality or cell death. Within the macrophage cluster, GFP^⁺^ maternal cells had a lower median mitochondrial transcript fraction than control pup macrophages (0.20% versus 0.66%; p = 1.3 × 10^⁻^□□), and a smaller proportion of cells contained >5% mitochondrial transcripts (0.7% versus 2.4%). Maternal macrophages had a median of 3,220 detected genes per cell. A score comprising 14 canonical apoptosis and cell-death genes was lower in maternal than control macrophages (p = 3.3 × 10^⁻^□), whereas a housekeeping and translational-machinery score did not differ between groups (p = 0.64; **Fig. S5C–E**). *Cdkn1a* was expressed in a higher proportion of maternal macrophages than control cells (63% versus 43%). Because the Flex probe panel provides limited mitochondrial coverage, the absolute mitochondrial transcript fractions are low; nevertheless, the same measurement was applied to both groups.

We next assessed whether the transcriptional differences between maternal and control macrophages were reproducible across the two biological replicates, which originated from independent litters collected five months apart. Differential expressions were analyzed separately for each replicate. Gene-level effect sizes were highly concordant between replicates (Pearson r = 0.948; Spearman ρ = 0.939 across 8,311 genes). Among the 1,356 genes reaching significance in either replicate, all showed the same direction of change in both replicates, and 969 genes were significant in both independently (**Fig. S5F**).

Chromosomal sex provided an additional population-level measure of cell composition. Cells sorted from pups nursed by GFP^⁺^ dams contained a lower proportion of Y-chromosome-transcript-positive cells than cells sorted from pups nursed by WT dams (13.2% versus 24.1%). Because the control libraries were sequenced more deeply, we tested whether this difference could be attributable to sequencing depth. After adjustment for sequencing depth, the difference remained significant (odds ratio = 0.50, p = 1.9 × 10⁻□), whereas sequencing depth itself was not significant. The difference was also maintained after downsampling all cells to a common depth of 1,500 UMIs (6.2% versus 9.3%, p = 0.027), across every quintile of sequencing depth, and when analysis was restricted to the macrophage cluster (4.0% versus 6.8%, p = 0.029; **Fig. S5G, H**).

### Flow cytometry and GFP mRNA analysis independently verify the presence of maternal cells in circulation

To independently validate the presence and identity of circulating maternal cells, we established a separate cohort of WT pups nursed by either WT or GFP^⁺^ dams and analyzed fresh blood samples by flow cytometry. The gating strategy consisted of sequential identification of total cells, viable cells using Zombie Violet, GFP^⁺^ cells, and CD45^⁺^CD11b^⁺^ myeloid cells (**Fig. S6**). Among pups nursed by GFP^⁺^ dams, 70.5±11.5% of total blood cells were viable (**Fig. 4A**). Within the GFP^⁺^ population, 39.9 ± 13.3% of cells were viable, and this population was subsequently used for RT-qPCR analysis. GFP^⁺^ cells constituted 0.21±0.05% of the viable cell population (**Fig. 4B**) and CD45 CD11b myeloid cells accounted for 76.3±2.3% of GFP cells (**Fig. 4C**). Importantly, these measurements were performed on freshly collected, unfixed samples and therefore provide an independent assessment of GFP^⁺^ cell recovery and viability, rather than a direct measure of viability in the fixed samples used for single-cell RNA sequencing. To further assess GFP expression at the transcriptional level, RNA isolated from the analyzed blood samples was subjected to RT-qPCR. GFP transcripts were detected in sorted cells from pups nursed by GFP dams, with a mean fold change of 10.7 relative to WT-dam controls (**Fig. 4D**; n = 2 per group). Inter-animal variability was substantial, reflecting the rarity of circulating maternal cells and the consequent variation in sorted cell yield.

**Figure 4.**
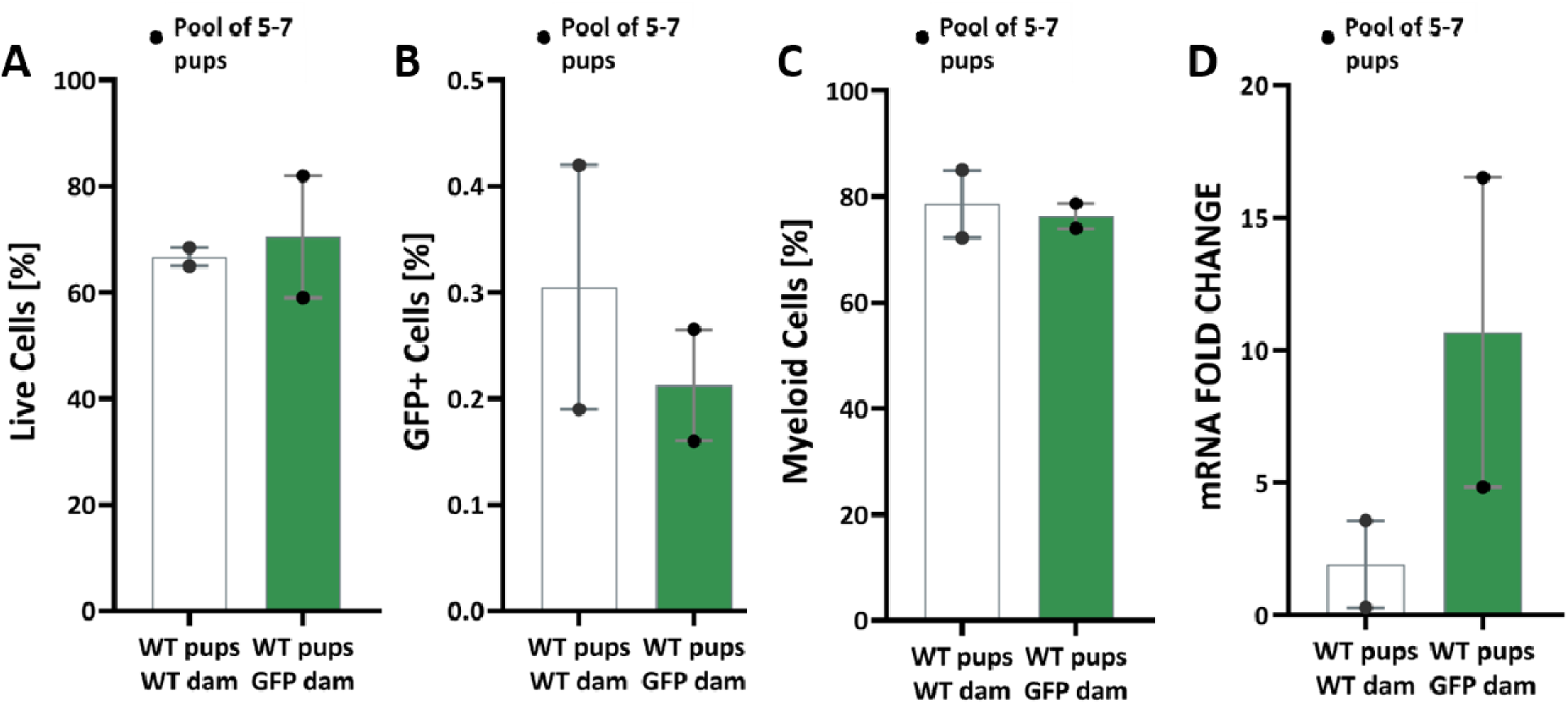
Circulating maternal cells are viable, predominantly myeloid, and contain detectable GFP transcript. **(A)** Percentage of live cells detected by flow cytometry in peripheral blood of WT pups nursed by WT dams or GFP^⁺^ dams. **(B)** Percentage of GFP^⁺^ cells among live cells in peripheral blood of WT pups nursed by WT dams or GFP^⁺^ dams, indicating the presence of maternally transferred cells. **(C)** Percentage of myeloid cells (CD45^⁺^CD11b^⁺^) among the analyzed cell population in both groups. **(D)** GFP mRNA expression quantified by qPCR for GFP^⁺^CD45^⁺^CD11b^⁺^ myeloid cells from pups nursed by GFP^⁺^ dams and GFP^⁺^ cells from pups nursed by WT dams to obtain sufficient cell numbers, presented as fold change relative to WT pups nursed by WT dams. Results are shown as mean ± SEM; n =2 independent litters per group.

The detection of GFP transcripts is nonetheless informative independently of the group comparison: GFP mRNA cannot be acquired through phagocytic uptake of GFP protein, or of extracellular vesicles carrying it, and its presence in sorted GFP cells therefore indicate that at least a fraction of these cells actively transcribes the maternal transgene.

### Maternal cells home to neonatal injury sites and adopt a myeloid phenotype

To determine whether the transcriptional signature of transferred maternal cells corresponds to functional migratory behavior *in vivo*, we employed a neonatal skin-injury model. WT pups cross-fostered to GFP^+^ transgenic dams from birth were nursed for nine days, after which a small dorsal skin incision was introduced. Pups continued nursing for an additional two days before tissue collection and analysis (**Fig. 5A**). Immunofluorescence imaging of skin sections stained for GFP and DAPI revealed abundant GFP^⁺^ maternal cells localized to wound sites in WT pups nursed by GFP^⁺^ dams, across three independent litters (**Fig. 5B**). In contrast, GFP□ cells were not detected above the background threshold defined by WT-nursed control sections, either in the injured skin of WT pups nursed by WT dams or in the healthy skin of WT pups nursed by GFP dams, indicating that maternal cell accumulation at wound sites is injury-dependent rather than a consequence of passive tissue dissemination. Quantitative analysis confirmed that GFP^⁺^ cells comprised 11±3% of DAPI-defined cells within the dermal wound region, corresponding to approximately 1,250±600 GFP^⁺^ cells per analyzed region (**Fig. 5C, D**). Both the percentage and absolute number of GFP^⁺^ cells were significantly elevated in injured pups nursed by GFP^⁺^ dams compared with both control groups.

**Figure 5.**
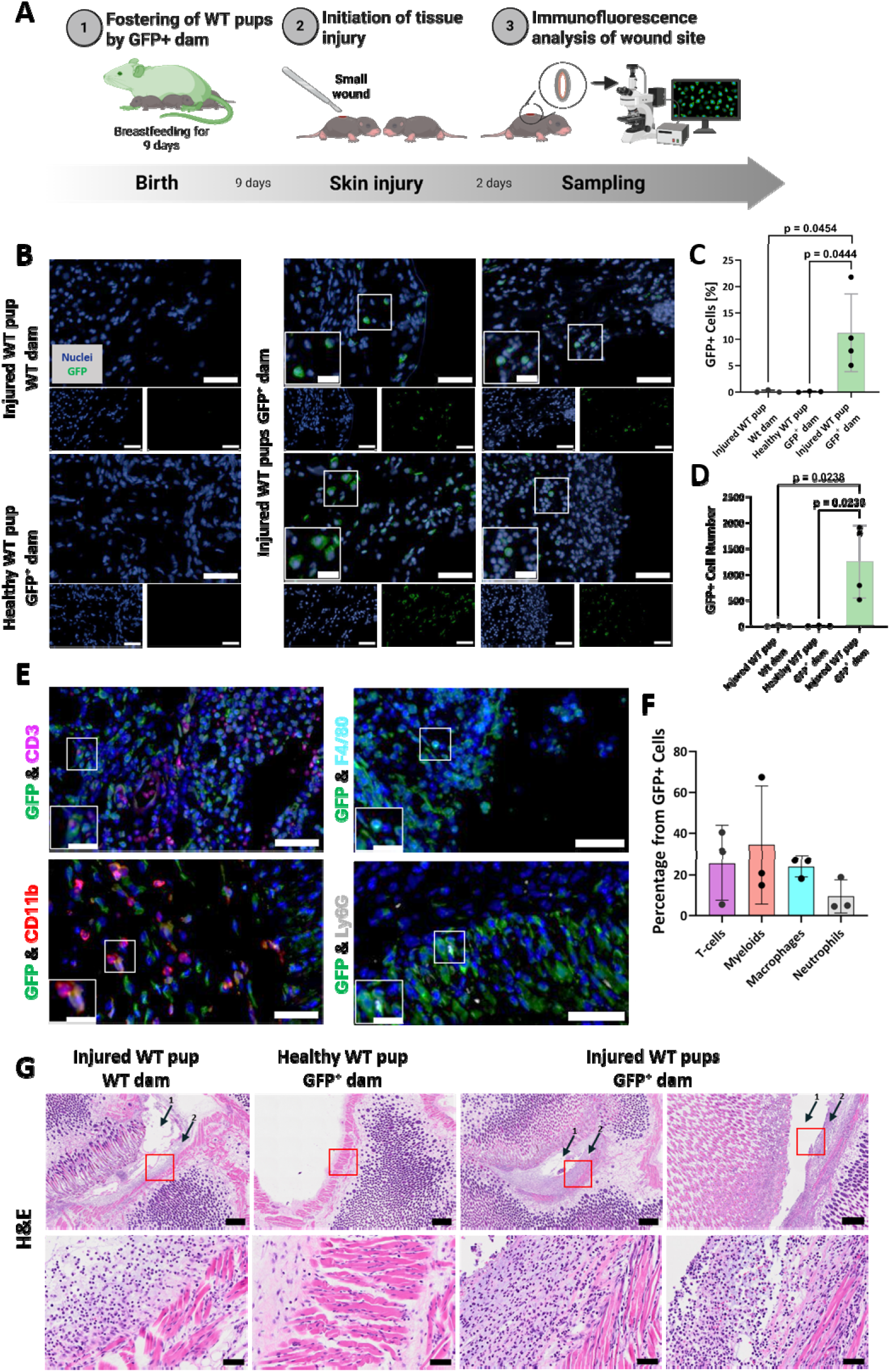
Breastmilk-derived maternal cells are detected at sites of neonatal tissue injury. **(A)** Experimental schematic of the neonatal skin-injury model. WT pups were cross fostered to GFP^⁺^ transgenic dams immediately after birth and nursed for 9 days. A small dorsal skin wound was then induced, followed by continued nursing and subsequent collection of injured tissue for histological and immunofluorescence analysis to assess maternal cell homing. **(B)** Representative immunofluorescence images of skin sections stained for nuclei (DAPI, blue) and GFP (green), imaged using an Olympus Slideview VS200 scanner. GFP^⁺^ cells are detected in injured skin of WT pups nursed by GFP^⁺^ dams but are absent from injured skin of WT pups nursed by WT dams and from healthy skin of WT pups nursed by GFP^⁺^ dams. Scale bars: 50 µm (overview); 20 µm (insets). Images were adjusted by +20 brightness and −40 contrast for display. **(C)** Percentage and **(D)** absolute number of GFP^⁺^ cells per field quantified across groups. **(E)** Representative immunofluorescence images of injured skin from WT pups nursed by GFP^⁺^ dams, co-stained for GFP (green) and CD3 (magenta, T cells), CD11b (red, myeloid cells), F4/80 (cyan, macrophages), or Ly6G (yellow, neutrophils). Insets highlight co-localization of GFP^⁺^ maternal cells with each marker. Scale bars: 50 µm (overview); 20 µm (insets). Images were adjusted by +20 brightness and −40 contrast for display. **(F)** Quantification of the immune cell composition among GFP^⁺^ maternal cells at the wound site, expressed as percentage of total GFP^⁺^ cells positive for each lineage marker (T cells, myeloids, macrophages, neutrophils). **(G)** Representative hematoxylin and eosin (H&E)-stained sections of injured skin from WT pups nursed by WT dams, healthy skin from WT pups nursed by GFP^⁺^ dams, and injured skin from WT pups nursed by GFP^⁺^ dams (two independent sections shown). Red boxes indicate regions shown at higher magnification below. Arrows mark the wound site and the underlying dermis layer. Injured tissues display characteristic inflammatory infiltration and disrupted tissue architecture, whereas healthy skin shows intact morphology. Scale bars: 500 µm (top rows); 50 µm (bottom rows). Data are presented as mean ± SEM. One-way ANOVA followed by Tukey’s multiple comparison test; P ≤ 0.05 considered statistically significant.

To characterize the immune identity of the recruited maternal cells, we performed multi-marker immunofluorescence co-staining of GFP^⁺^ cells with lineage-specific markers including CD3 (T cells), CD11b (myeloid cells), F4/80 (macrophages), and Ly6G (neutrophils) (**Fig. 5E**). GFP^⁺^ maternal cells were detected co-expressing each of these markers within the wound dermis, with insets confirming co-localization at the single-cell level. Quantification of the immune composition among GFP^⁺^ cells revealed that myeloid cells and macrophages represented the most abundant transferred populations (34±17% and 24±3%, respectively), followed by T cells (25±10%) and neutrophils (9±5%) (**Fig. 5F**). Data are presented as mean ± SEM. Notably, a substantial fraction of GFP cells co-expressed the T cell marker CD3. Because T cells do not internalise whole cells or large cellular fragments, GFP□CD3□ events cannot be accounted for by phagocytic uptake of maternal material by host cells and therefore provide independent support for the presence of intact transferred maternal cells. This phenotypic distribution is consistent with the monocyte-macrophage transcriptional identity identified by scRNA-seq and with the chemokine-responsive gene expression profile observed in circulating maternal cells.

Histological analysis of H&E-stained sections confirmed the integrity and morphological characteristics of each experimental condition (**Fig. 5G**). Injured skin from WT pups nursed by WT dams displayed characteristic wound architecture with dense inflammatory infiltration in the dermis, as indicated by arrows marking the wound site and underlying dermal layer. Healthy skin from WT pups nursed by GFP^⁺^ dams showed intact epidermal and dermal morphology without evidence of inflammatory disruption. Injured skin from WT pups nursed by GFP^⁺^ dams exhibited comparable wound morphology with similar prominent inflammatory infiltration, confirming equivalent tissue injury across groups. Red-boxed regions are shown at higher magnification in the lower panels to illustrate the cellular composition of the dermal compartment. Collectively, these data demonstrate that breastmilk-derived maternal cells selectively home to sites of neonatal tissue injury in vivo, adopt a predominantly myeloid phenotype within the wound microenvironment, and do so in a manner consistent with active chemotactic recruitment rather than passive tissue seeding.

## Discussion

Maternal–offspring cellular exchange is well established during gestation. Yet, the extent to which intact maternal cells are transferred to the offspring after birth, persist systemically, and acquire defined identities within the offspring has remained unsolved^23,42^. We provide evidence consistent with breastfeeding constituting a postnatal route for maternal cell transfer into the neonate. This study demonstrated that maternal cells transferred to neonates through breastfeeding can survive gastrointestinal transit, access the circulation, localize to peripheral organs, and accumulate at inflammatory sites. By combining *in vivo* tracking with single-cell transcriptomics, we further define the cellular composition and functional state of these transferred cells, supporting the concept that breastfeeding may serve as an active biological pathway for maternal cellular input into the neonate, independent of pregnancy, and complementing current understanding of breastmilk-mediated communication beyond soluble components such as antibodies and cytokines^43,44^.

Our findings extend and refine previous observations of postnatal maternal cell transfer. Earlier studies have suggested that maternal cells can be transmitted to offspring both during gestation and through breastfeeding. Zhou et al.^16^ described two independent routes of maternal cell transmission-placental transfer during pregnancy and breastfeeding after birth-establishing the conceptual framework for postnatal maternal cell trafficking. Subsequent work demonstrated that maternal cytotoxic T lymphocytes derived from breastmilk can localize to Peyer’s patches in nursing offspring^15^, and that breastmilk cell trafficking may contribute to immune system maturation through microchimerism-mediated mechanisms^45^. In addition, Darby et al.^19^ showed that pathogen-experienced maternal T cells transferred through breastfeeding can confer long-term antigen-specific protection in offspring. More recently, Doerfler et al.^32^ reported that maternal milk cell components can also be taken up by neonatal macrophages through extracellular vesicle–mediated transfer, suggesting that cell-free material exchange may represent a complementary pathway. Our results build on these studies by demonstrating that intact maternal immune cells can enter neonatal circulation after gastrointestinal passage and by providing single-cell transcriptomic evidence that these transferred cells comprise multiple hematopoietic lineages, with macrophages representing the dominant population. The transcriptional profiles we observe, characterized by inflammatory priming coupled with metabolic restraint, further indicate that transferred maternal cells adopt a distinctive stress-adapted state within the neonatal environment. An important strength of this model is that prenatal transfer was excluded by design. WT pups were delivered by WT mothers and only subsequently fostered by GFP-transgenic dams, ensuring that GFP^⁺^ cells identified in recipient tissues originated from postnatal maternal transfer through lactation rather than transplacental maternal microchimerism. Together, these findings provide molecular evidence supporting the transfer of viable maternal immune cells through breastfeeding while remaining compatible with the possibility that extracellular vesicle–mediated material transfer occurs in parallel.

A key observation in this study is that breastmilk-derived maternal cells were detected in neonatal blood, with lower representation in the liver and lungs. Although paraffin processing precludes visualization of native GFP fluorescence, antibody-based detection provided the sensitivity required to identify the very small population of transferred maternal cells in recipient tissues. This distribution argues against indiscriminate tissue seeding and instead supports a model in which postnatal maternal cell trafficking is constrained by tissue accessibility and local cues. The absence of detectable maternal cells in the brain, kidney, and spleen further suggests that entry into neonatal tissues is selective rather than stochastic. Whether this pattern reflects physical barriers, tissue-specific vascular architecture, differential retention, or rapid clearance in some compartments remains to be determined. Nonetheless, these data establish that maternal cell transfer can be generated after birth, independently of pregnancy, and that systemic dissemination is not limited to the intestinal mucosa.

Our intestinal imaging is consistent with earlier reports demonstrating that breastmilk-derived maternal immune cells are located to the neonatal intestinal mucosa and Peyer’s patches following nursing^14,15^. Previous studies primarily focused on documenting the presence of maternal cells within the gastrointestinal tract, supporting the intestine as an important site of maternal– offspring cellular interactions. Our findings reproduce these observations in an independent foster-nursing model while extending them by showing that maternal cells are subsequently detected in the circulation, exhibiting a distinct transcriptional profile dominated by stress-adapted macrophages, and can localize to distant inflammatory sites. Together, these results suggest that the intestine represents the initial portal of maternal cell entry, whereas systemic dissemination may enable functions beyond the gastrointestinal tract.

A central advance of our work is the molecular characterization of transferred maternal cells after gastrointestinal passage. Single-cell RNA sequencing revealed a heterogeneous maternal cell pool encompassing multiple hematopoietic lineages, indicating that milk can deliver diverse immune cell types into the neonatal circulation. Within this pool, the monocyte/macrophage compartment was particularly informative: transferred maternal macrophages formed a transcriptionally distinct population relative to endogenous pup macrophages, segregating clearly in low-dimensional embedding and exhibiting coordinated differences in gene expression. This separation was not subtle, suggesting that transferred maternal cells retain lineage identity while adopting a unique state shaped by transit and/or adaptation to the neonatal host environment.

An important consideration in interpreting the transcriptomic data is whether the GFP^⁺^ cells represent cells of maternal origin or neonatal cells that have acquired maternal GFP-containing material. We addressed this question using several complementary approaches. Quantitative PCR detected GFP transcripts in RNA isolated from independently sorted GFP^⁺^ cells, supporting the presence of cells actively transcribing the maternal transgene, although the magnitude of the difference relative to WT controls varied between animals and did not reach statistical significance. Because GFP mRNA cannot be acquired through phagocytosis of GFP protein alone, these findings provide supporting evidence for the presence of maternal cells rather than simple transfer of GFP protein. Consistent with this, EGFP transcripts were detected exclusively in cells from pups nursed by GFP^⁺^ dams and were absent from all 1,390 cells analyzed from WT-nursed controls. However, EGFP detection was sparse, and individual EGFP^⁺^ cells could not be assigned unequivocally to maternal origin; some also contained Y-chromosome transcripts. We therefore interpret EGFP detection as evidence that transgene-transcribing cells are present in the experimental population rather than as a definitive cell-by-cell marker of maternal origin.

Additional population-level evidence was provided by the reduced representation of Y-chromosome-transcript-positive cells in the experimental samples compared with controls. This difference persisted after adjustment for sequencing depth, after downsampling to a common sequencing depth, and when analysis was restricted to the macrophage population. Although this pattern is consistent with the contribution of female maternal cells to the sorted population, it should not be interpreted as a quantitative estimate of maternal-cell frequency. The analysis was based on a single control litter, and the sex composition of the pooled litters was not recorded, preventing retrospective verification that the experimental and control groups had comparable sex ratios. Nevertheless, the magnitude of the estimated contribution was comparable to the independent GFP^⁺^ frequencies obtained by flow cytometry, providing convergent evidence from distinct experimental approaches. Together with the qPCR results, EGFP transcript detection, the presence of viable GFP^⁺^ cells, and the coherent transcriptional divergence of GFP^⁺^ macrophages from endogenous pup macrophages, these observations support the interpretation that the single-cell transcriptional profiles reflect transferred maternal cells rather than neonatal cells that have passively acquired maternal GFP protein or cellular material.

The possibility that the observed transcriptional state reflected preferential recovery of dying cells was also examined. Maternal macrophages showed lower mitochondrial transcript fractions and lower expression of a canonical apoptosis and cell-death gene set than control macrophages, while housekeeping and translational-machinery scores were comparable between groups. These findings argue against poor cell quality or cell death as the primary explanation for the maternal macrophage transcriptional state. Finally, the strong concordance of differential expression across the two independently processed biological replicates (Pearson r = 0.948; Spearman ρ = 0.939) provides additional support for the reproducibility of the transcriptional phenotype. Although the limited number of biological replicates warrants caution in generalizing these findings, the consistent direction of differential expression across replicates strengthens the conclusion that the observed macrophage state represents a reproducible biological signal rather than a replicate-specific effect.

The transcriptional program of transferred maternal macrophages was notable for the combination of inflammatory priming with metabolic restraint. Enrichment of AP-1-associated activity and, TNFa–NF-κB signaling, together with TLR4 signaling pathways that include MyD88-independent components, suggests exposure to inflammatory or microbial-associated cues^46^. In parallel, maternal macrophages showed marked suppression of ribosomal and translational programs, reduced mitochondrial-associated signatures, and downregulation of pathways involved in lipid homeostasis and RNA processing. This coupling of immune activation signatures with broad suppression of biosynthetic and energetic pathways is consistent with a stress-adapted state, in which cells prioritize persistence under challenging conditions over proliferation and growth^47,48^. Supporting this interpretation, maternal cells were underrepresented in the cycling monocyte/macrophage cluster enriched for proliferative markers, arguing against extensive post-transfer expansion at this time point.

Notably, the transcriptional features observed in maternal transferred cells were reflected in their behavior *in vivo*. In a neonatal skin injury model, maternal GFP^⁺^ cells were detected at wound sites and were absent from healthy skin, consistent with selective recruitment rather than passive dissemination. This spatial restriction argues against direct cutaneous transfer as a route of maternal cell entry, as such a mechanism would be expected to yield detectable maternal cells in uninjured skin. Instead, the data are more consistent with a model in which maternal cells first traverse the neonatal gastrointestinal tract, access the circulation, and are subsequently recruited to sites of inflammation in response to local cues, although passive trapping within inflamed vasculature cannot be fully excluded. The detection of CD11b^⁺^ maternal cells within injured tissue is consistent with the myeloid identity and chemokine-responsive signatures identified by single-cell analysis, although a direct functional link between transcriptional state and migratory behavior remains to be established. Importantly, although these findings show that breastmilk-derived maternal cells can localize to inflamed tissue, they do not establish active recruitment, a causal contribution to wound healing, inflammatory regulation, or tissue remodeling. Defining whether transferred maternal cells actively modulate neonatal immune responses or tissue repair, and whether their impact is beneficial, neutral, or context-dependent-will require targeted perturbation approaches and functional readouts.

These findings raise several broader implications for understanding neonatal immune biology and maternal–offspring crosstalk. Early postnatal life is characterized by rapid microbial colonization, barrier maturation, and immune adaptation^49,50^. Maternal immune cells transferred through breastfeeding may represent an additional layer of maternal influence during this critical developmental window, complementing antibodies, cytokines, metabolites, and microbial exposures present in milk^51^. While the stress-adapted yet immunologically poised transcriptional state of transferred maternal macrophages is consistent with a capacity to sense inflammatory environments, whether these cells actively interact with neonatal immune networks, and whether such interactions shape immune education, tolerance, susceptibility to infection, or later-life inflammatory trajectories, remains speculative and will require direct functional investigation.

The pathophysiological relevance of these observations deserves explicit consideration. The neonatal immune system is characterized by quantitative and qualitative deficiencies in myeloid function: neonatal monocytes and macrophages exhibit blunted pro-inflammatory cytokine production, reduced phagocytic efficiency, and limited capacity for oxidative burst compared with their adult counterparts, deficits that are believed to underlie neonatal susceptibility to invasive bacterial and fungal infections. Maternal macrophages transferred through breastfeeding, which our data show is transcriptionally primed with inflammatory and TLR4 signaling signatures, may have the potential to contribute to innate immune responses at precisely the developmental stage when endogenous myeloid responses are most limited. Consistent with this concept, the recruitment of maternal GFP^⁺^ cells to sites of neonatal skin injury, and their co-expression of the myeloid marker CD11b, suggests that these cells may participate in tissue responses to damage-associated signals, not merely passively distribute through neonatal tissues. Beyond acute infection responses, maternal macrophage transfer could contribute to neonatal mucosal immune homeostasis: maternal cells traversing the intestinal barrier are exposed to luminal microbial antigens before systemic entry, potentially acting as antigen-presenting intermediaries that shape early neonatal immune tolerance and microbial recognition thresholds. Conversely, if maternal cells transferred during breastfeeding are dysregulated, for example carrying inflammatory programmes associated with maternal autoimmune disease, systemic infection, or metabolic stress they could represent a cellular vehicle for adverse immune programming in the offspring, potentially contributing to neonatal inflammatory conditions or to longer-term immune dysregulation. While the current study cannot resolve whether the transferred maternal cells observed here are functionally protective, neutral, or context-dependent in their effects, the evidence that they survive gastrointestinal transit, adopt a stress-adapted yet immunologically primed state, and home to sites of tissue injury collectively supports a model in which postnatal maternal cell transfer represents a biologically meaningful pathway, one that warrants direct functional investigation using loss-of-function approaches, infection models, and longitudinal immune phenotyping in both murine and ultimately human neonatal systems.

Our study also has limitations that define clear next steps. First, the duration of persistence beyond the nursing period and the fate of transferred maternal cells within tissues were not resolved here. Longitudinal tracking will be needed to determine whether these cells are transient passengers, short-term responders, or longer-lived chimeric residents. Second, our single-cell profiling focused on circulating maternal cells; whether tissue-localized maternal cells share the same transcriptional states, further differentiate within organs, or acquire tissue-imprinted programs warrants direct analysis. Third, while our data support selective recruitment to injured tissue, the molecular determinants of trafficking-such as specific chemokine receptors–ligand axes- and the factors governing tissue restriction remain to be defined. Fourth, although GFP transcripts were detected in sorted GFP cells, this orthogonal validation is constrained by the small number of cells recoverable per animal and the resulting inter-animal variability; genomic confirmation of donor origin, for example by donor-specific genotyping of sorted cells, would further strengthen lineage assignment. Fifth, because milk composition and immune-cell content can vary with lactation stage, maternal immune status, and environmental exposures, it will be important to establish how generalizable these findings are across conditions and whether maternal inflammation or infection alters the quantity or state of transferred cells. Sixth, it is important to note that all conclusions presented here are derived from a murine model using controlled foster-nursing and injury paradigms. Although this system enables precise dissection of postnatal maternal cell transfer and functional behavior, important species-specific differences exist in lactation biology, neonatal immune development, gastrointestinal permeability, and maternal–offspring immunological crosstalk. Therefore, caution should be exercised when extrapolating these findings directly to humans. Whether breastmilk-derived maternal immune cells similarly survive gastrointestinal transit, enter the circulation, persist in tissues, and functionally integrate within the human neonate remains to be determined and will require dedicated translational and clinical investigation.

In summary, we identify breastfeeding as a postnatal route of maternal cellular transfer, demonstrate that breastmilk-derived maternal immune cells can access neonatal circulation and select organs, and show that transferred maternal macrophages adopt a distinctive transcriptional program characterized by inflammatory readiness coupled to metabolic suppression and stress adaptation. The recruitment of maternal cells to neonatal injury sites provides *in vivo* evidence consistent with responsiveness to inflammatory cues within the offspring. Together, these results contribute to emerging evidence that maternal– neonatal crosstalk may include viable cellular transfer after birth and highlight the need for future studies to define the persistence, tissue fate, and physiological consequences of breastmilk-derived maternal cell transfer.

## Methods

### Foster-nursing *in vivo* model

C57BL/6-Tg(UBC-GFP)30Scha/J mice^38^ (The Jackson Laboratory) and WT C57BL/6 mice (age 6 weeks old) were bred in parallel to obtain synchronized pregnancies. To exclude prenatal transfer of maternal GFP^⁺^ cells, all experimental pups were born to wild-type dams and were adopted by GFP-transgenic foster dams immediately after birth. Consequently, any GFP^⁺^ cells detected in recipient pups originated exclusively from postnatal transfer during lactation. A single GFP^+^ pup was left with each dam to improve acceptance. WT fostered pups were collected after eighteen days of nursing for downstream analysis. Whole blood was collected into EDTA tubes (BD), and organs were harvested. For the injury model, WT pups fostered for nine days were subjected to a small dorsal skin incision. After two days, pups were euthanized and wound tissues were collected for analysis. All procedures were approved by the institutional animal ethics committee (protocol numbers IL-1521022, IL-0960725 and IL-1521124).

### Blood preparation

Fresh murine blood was diluted in 1 mL of RBC lysis buffer (Sigma-Aldrich), incubated at room temperature with gentle agitation, and centrifuged at 500 × g for 10 min. Pelleted cells were resuspended in 1 mL RBC lysis buffer, centrifuged again, and finally resuspended in PBS.

### Organ dissociation

Organs were minced into small fragments and digested in PBS containing Collagenase type IV (2.5 mg/mL), Collagenase type III (2.5 mg/mL), and Hyaluronidase type I-S (2.5 mg/mL) at 37 °C for 45 minutes with agitation every 15 minutes. Cell suspensions were passed through a 70-µm nylon strainer and centrifuged at 200×g for 3 minutes, followed by centrifugation at 400×g for 3 minutes. Red blood cell lysis was performed as described for blood samples.

### Flow cytometry analysis

For flow cytometry analysis, approximately 1×10 cells were stained with Zombie Violet™ live/dead dye (BioLegend) diluted 1:500, incubated for 20 minutes at room temperature in the dark, washed with PBS containing 2% FBS, and analyzed for GFP expression using a BD LSR-II flow cytometer (LS&E Infrastructure Center, Technion). Data was analyzed using FlowJo software (BD Biosciences). The gating strategy consisted of sequential exclusion of debris based on forward- and side-scatter characteristics, followed by singlet discrimination using FSC-A versus FSC-H to remove cell aggregates. Viable cells were subsequently identified by excluding Zombie Violet-positive events, and GFP fluorescence was analyzed within the live singlet population. GFP-positive cells were quantified as a percentage of the total live singlet population and used for subsequent analyses.

### Cell sorting

Peripheral blood was collected from WT pups nursed by GFP-expressing dams (experimental group) and age-matched WT pups nursed by WT dams (control group). For single-cell RNA sequencing, blood samples were processed according to the Chromium Fixed RNA Profiling protocol (10x Genomics). Briefly, cells were fixed overnight at 20 °C using 4% formaldehyde and Fix & Perm buffer (10x Genomics). Following fixation, cells were centrifuged at 850 × g for 5 min and resuspended in 500 µL chilled Quenching Buffer (10x Genomics). Cells were sorted at 4 °C using a Bigfoot cell sorter (Invitrogen; LS&E Infrastructure Center, Technion) and collected into Quenching Buffer. Cellular events were identified based on forward- and side-scatter characteristics, and debris was excluded prior fluorescence-based sorting. GFP^⁺^ cells were collected from the experimental group (same instrument settings and gating criteria were applied for the control group). As cells were fixed prior to sorting, viability discrimination was not performed. No singlet discrimination gate or additional side-scatter exclusion was applied to preserve the broadest representation of circulating immune cell populations. Thus, experimental and control samples underwent identical fixation, sorting, and collection procedures prior to library preparation and single-cell RNA sequencing.

For qPCR analysis, unfixed cells were stained with Zombie Violet™ live/dead dye (BioLegend; 1:500 dilution), for 20 minutes at room temperature in the dark, washed with PBS containing 2% FBS, and stained with 2ug/ml PE-conjugated anti-CD45-(30-F11) (BLG103105, BioLegend) and 5ug/ml Alexa Fluor 647-conjugated anti-CD11b-(M1/70) (BLG101218, BioLegend) antibodies for 30 minutes at 4 °C in the dark. Cells were analyzed by flow cytometry, and debris was excluded based on forward- and side-scatter characteristics. Viable cells were identified by exclusion of Zombie Violet-positive events, followed by identification of GFP^⁺^ cells within the live singlet population. CD45^⁺^CD11b^⁺^ cells within the GFP^⁺^ population were classified as myeloid cells. For downstream RNA extraction and qPCR analysis, GFP^⁺^ cells from WT pups nursed by WT dams and GFP^⁺^CD45^⁺^CD11b^⁺^ myeloid cells from WT pups nursed by GFP-expressing dams were sorted with a minimum target of 1,000 cells. Sorting continued until the sample was exhausted. Flow cytometry data were analyzed using FlowJo software (BD Biosciences).

### Single cell RNA sequencing

All samples were processed using a uniform workflow to minimize technical variability using the 10x Genomics Chromium Fixed RNA Profiling (Flex) platform, according to the manufacturer’s protocol. Because breastmilk-derived maternal cells are rare and highly fragile following fluorescence-activated cell sorting, cells were fixed prior sorting to stabilize cellular RNA and minimize stress-induced transcriptional artifacts prior to droplet encapsulation. Fixed cells were subsequently loaded onto the Chromium X instrument to generate gel bead–in–emulsion (GEM) droplets, targeting the capture of approximately 5,000 cells per channel. Following cDNA amplification, sequencing libraries were generated and sequenced on an Illumina NextSeq 2000 platform (Azrieli Technion Genomics Center) in accordance with the manufacturer’s instructions. Libraries were prepared using the Chromium Mouse Transcriptome Probe Set v1.0.1 supplemented with three custom probes targeting EGFP (EGFP-1, EGFP-2 and EGFP-3), enabling direct detection of transgene transcripts in the single-cell data. Counts from the three EGFP probes were summed into a single EGFP feature prior to analysis.

### Data processing and quality control

Raw sequencing data were processed using Cell Ranger (10x Genomics). Count matrices were analyzed in R using Seurat^52^. Across samples, the median genes per cell ranged from 1,643 to 4,136 and median UMI counts per cell ranged from 3,207 to 12,324. The three samples (two GFP^⁺^ replicates, one control) were merged without integration. Doublets were identified using scDblFinder^53^ (version 1.24.0) and removed from downstream analyses. EGFP-transcript-positive cells were defined as cells containing at least one unique EGFP molecular identifier (UMI), and the frequency of EGFP-positive cells was compared between groups using Fisher’s exact test. Apoptosis and housekeeping scores were calculated as the mean normalized expression across fourteen canonical cell-death genes (Casp3, Casp7, Casp8, Casp9, Bax, Bak1, Bid, Apaf1, Pmaip1, Bbc3, Trp53, Cdkn1a, Aifm1 and Endog) and nine housekeeping and translational-machinery genes, respectively. Scores were compared between maternal and control macrophages using the Wilcoxon rank-sum test. Biological replicate concordance was assessed by performing differential expression analysis between maternal and control macrophages separately for each GFP^⁺^ replicate and correlating the resulting log□ fold changes. Chromosomal sex was assessed based on expression of the Y-linked genes Ddx3y, Eif2s3y, Uty and Kdm5d, all represented in the probe panel. A cell was classified as Y-positive if at least one UMI was mapped to any of these genes. Because sequencing depth differed between groups, the comparison was repeated using three complementary approaches: logistic-regression adjustment for log□□ total UMI counts, downsampling of all cells to a common depth of 1,500 UMIs per cell, and stratification by sequencing-depth quintile.

### Clustering and cell type annotation

Following log-normalization and identification of the top 2,000 highly variable features, principal component analysis was performed, and the first 15 principal components were used for UMAP dimensionality reduction and clustering (resolution = 0.1). Cell types were annotated using SingleR^40^ with ImmGenData and MouseRNAseqData reference datasets, supplemented by manual annotation based on canonical marker genes.

### Macrophage subset analysis and differential expression

Cluster 0, which comprised the largest macrophage population and exhibited clear segregation between GFP^⁺^ and GFP^⁻^ cells, was subset for focused analysis. Differential gene expression between GFP^⁺^ maternal macrophages and GFP^⁻^ control macrophages was assessed using the Wilcoxon rank-sum test with genes detected in ≥10% of cells. Genes with adjusted p-value < 0.05 (FDR correction) and absolute log□ fold-change > 1 were considered significantly differentially expressed.

### Pathway enrichment analysis

Gene set enrichment was performed using Ucell (version 2.14.0) with gene sets from MSigDB, including Hallmark, Gene Ontology, Reactome, and immunologic signatures (**Table S1**). Pathway scores were calculated for individual cells (maxRank = 1,000) and z-score normalization within each pathway. Statistical comparison between GFP^⁺^ and control cells used the Wilcoxon rank-sum test with effect sizes quantified by Cohen’s d. Pathways with FDR-adjusted p-value < 0.05 were considered significantly enriched.

### Comparative analysis to lactation-associated macrophages

To assess whether our identified maternal macrophages correspond to lactation-associated macrophages (liMacs), we performed an integrated single-cell transcriptomic analysis of our cells and a published liMac macrophage dataset ^41^. Only macrophage populations were included in this analysis. The liMac dataset comprised macrophage populations isolated from lactating mammary tissue, as described in the original study.

Single-cell datasets were normalized independently using SCTransform (Seurat v5) with default parameters. Integration features were selected across datasets, and shared anchors were identified using SCT-based integration. Datasets were integrated using the IntegrateData function, and dimensionality reduction was performed using principal component analysis followed by UMAP.

To enable quantitative comparison of lineage identity and transcriptional state, we defined four gene modules: 1) Macrophage core module - consisted of canonical macrophage lineage markers and was used to confirm shared macrophage identity across datasets; 2) liMac module - defined using the full gene set reported as enriched in liMacs relative to other macrophage subsets in the original liMac study; 3) GFP-up module - comprising genes upregulated in GFP^⁺^ macrophages relative to control macrophages; and, 4) GFP-down module -comprising genes downregulated in GFP^⁺^ macrophages. Module scores were calculated at the single-cell level using rank-based module scoring (UCell) applied to the RNA assay.

### Quantitative Real-Time RT-PCR

Total RNA was extracted from the sorted cells using TRIREAGENT (Sigma Aldrich) according to the manufacturer’s protocol. RNA was reverse transcribed using High-Capacity cDNA Reverse Transcription Kit (Applied Biosystems Inc.). qRT-PCR was performed using PowerTrack™ SYBR Green Master Mix (Thermo Fisher Scientific) on a QuantStudio 1 instrument (Applied Biosystems). Values were normalized to the RPL13a housekeeping gene. Primer sequences are listed below. Data are presented as mean fold change compared to WT control group (sorted cells from WT pups nursed by WT dam) using the 2^-ΔΔCT^ method.

Primer sequences used for qRT-PCR (mouse origin):

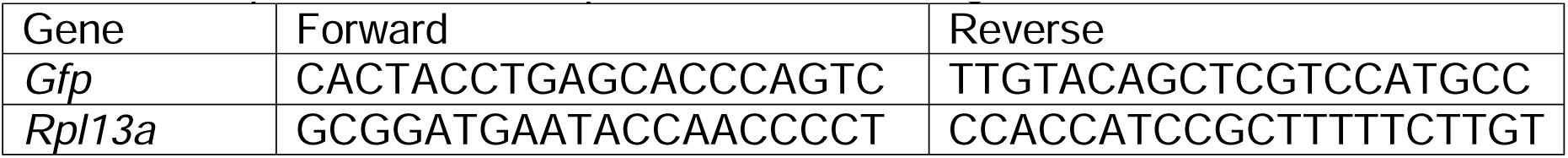

### Paraffin Sections preparation

Tissues were fixed in 4% paraformaldehyde, washed in PBS, dehydrated, and cleared using a Leica TP1020 tissue processor. Samples were embedded in Paraplast (Leica) using a Histocore Arcadia system and sectioned at 5 µm using a Leica RM2265 microtome. Sections were floated on a 37–39 °C water bath, mounted on adhesive slides (Leica X-tra), and dried overnight at 37 °C.

### Immuno-fluorescence and H&E staining

Sections were deparaffinized in xylene, rehydrated through graded ethanol, and subjected to antigen retrieval in citrate buffer (10 mM citric acid, 0.05% Tween-20, pH 6.0). After blocking (20% FBS, 0.2% Triton X-100), skin sections were incubated overnight with 12ug/mL anti-GFP polyclonal antibody (ab6673, Abcam) in 2% FBS, 0.2% Triton X-100, followed by a 1-h incubation with bovine anti-goat Alexa Fluor-647-conjugated secondary antibody (7ug/ml, Jackson ImmunoResearch) and 1:1000 diluted DAPI (Sigma). Additionally, two antibody panels were used for the skin sections. In addition to anti-GFP and DAPI as described above, one panel included 7ug/mL anti-Ly6G-[EPR22909-135] (ab238132, Abcam) and 10ug/mL anti-F4/80-[CI:A3-1] (ab6640, Abcam) antibodies, whereas the second included 5ug/mL anti-CD11b-[1250D] (MAB11242, R&D systems) and 5ug/mL anti-CD3-[17A2] (14-0032-82, eBioscience) antibodies followed by donkey anti-rabbit Cy3-conjugated (7ug/ml, Jackson ImmunoResearch) and donkey anti-rat Alexa Fluor-488-conjugated (7ug/ml, Jackson ImmunoResearch) secondary antibodies. The intestine sections stained with 12ug/mL anti-GFP (ab6673, Abcam), 2.5ug/ml anti-E-cadherin-[36] (BD610182, BD Biosciences) and 4.5ug/ml anti-CD31-[EPR17259] (ab124432, Abcam) primary antibodies for overnight at 4 °C in 2% FBS and 0.05% Tween-20 in PBS. Followed by 1-h incubation with bovine anti-goat Alexa Fluor-647-conjugated (7ug/ml, Jackson ImmunoResearch), donkey anti-mouse Alexa Fluor 488-conjugated (7ug/ml Jackson ImmunoResearch), and donkey anti-rabbit Cy3-conjugated (7ug/ml, Jackson ImmunoResearch) secondary antibodies and 1:1000 diluted DAPI (Sigma).

Tissues were processed as paraffin sections for histological evaluation and immunostained with an anti-GFP antibody rather than relying on endogenous GFP fluorescence. This approach was selected to maximize the sensitivity of detecting the rare transferred GFP-positive maternal cells while enabling thin (4– 5 μm) sections with excellent preservation of skin morphology, allowing accurate localization of individual cells within the epidermis and dermis. For H&E staining, sections were stained with hematoxylin and eosin and coverslipped using a Leica Histocore Spectra instrument.

### Image acquisition and quantification

Slides were scanned using an Olympus VS200 slide scanner (Olympus Corporation, Tokyo, Japan) with a 20x objective lens (0.325 μm/pixel resolution). The LED excitation and emission filter combinations used were: 385 nm excitation with 432/36 nm emission filter for DAPI, 555 nm excitation with 595/31 nm emission filter for Cy3, and 635 nm excitation with 680/42 nm emission filter for Cy5. For quantitative analysis, one tissue section was analyzed per wound, and one field encompassing the dermal area surrounding the wound was selected. Quantitative image analysis was performed on the original, unprocessed images using QuPath v0.6.0. Cells were identified using QuPath’s cell detection plugin based on DAPI staining, with a 5 µm cell expansion. A suitable threshold was then applied to the GFP detected channel, and the percentage of GFP-positive cells was calculated relative to the total number of cells in the selected dermal region. For multiple cell labeling, the threshold was applied for each fluorescent channel and the percentage of Ly6G, F4/80, CD11b or CD3-positive cells was calculated relative to the number of GFP-positive cells in the selected dermal region. For figure presentation only, skin representative fluorescence images were uniformly adjusted for brightness (+20) and contrast (−40), as indicated in the corresponding figure legends. The same adjustments were applied identically to all experimental groups within each figure, and no image-specific modifications were performed. These adjustments were used solely for visualization and were not used for quantitative analysis.

### Statistical analysis

Data are presented as mean ± standard error of the mean (SEM). Statistical significance was determined using one-way ANOVA with Tukey’s multiple comparison test for comparisons, and one-tailed unpaired Student’s t-test for comparisons between two independent groups. Pups from a single litter were pooled to obtain sufficient numbers of rare circulating maternal cells; each data point therefore represents one independently processed pooled sample, and independent litters were treated as biological replicates. The statistical test used, the experimental unit, and the number of independent biological replicates are stated in each figure legend. Analyses were performed using GraphPad Prism (GraphPad Software, La Jolla, CA), and p ≤ 0.05 was considered statistically significant.

## Acknowledgments

While preparing this work, we used ChatGPT 5.2 and Claude 1.4 to improve the readability and language of the manuscript. After using this tool, we reviewed and edited the content as needed and took full responsibility for the publication’s content. GFP^⁺^ mice were generously provided by Dr. Yitzhak Reizel, Department of Biotechnology and Food Engineering, Technion – Israel Institute of Technology. The *in vivo* experiments were performed with the support of Vita Zlobin, Yonatan Nisan and Dr. Amit Avrahami from the Technion Pre-Clinical Research Authority. Flow cytometry and cell sorting experiments were performed at The Life Science and Engineering (LS&E) Infrastructure Center at the Technion with the support of Aviv Lutaty and Yousef Mansour. Section preparation and histology staining were performed by Katren Sakran, Biomedical Core Facility (BCF) at the Technion. Imaging was performed by Maya Holdengreber, Melia Gurewitz and Adem Hind from the Biomedical Core Facility (BCF) at the Technion. The sequencing was performed with the support of the Azrieli Technion Genomics Center. The authors thank the Assaf Zinger laboratory team for their scientific and technical support.

## Author contributions

A.Z., S.N., and D.A conceived this study and designed the experimental strategy. S.N., T.G.L., S.E., and A.Z, performed the *in vivo* experiments, sample preparation, and flow cytometry analyses. C.Y.-B.U., M.B.M., and L.H.C. performed single-cell RNA sequencing experiments. A.A. and D.A. performed computational and bioinformatic analyses, including single-cell transcriptomic analysis and pathway enrichment. S.N., D.A., and A.Z., interpreted the data and developed the conceptual framework of the study. S.N., D.A., and A.Z. wrote the initial manuscript draft with input from all authors. All authors contributed to data discussion, manuscript editing, and approved the final version. D.A and A.Z. provided supervision, project administration, and resources. A.Z. secured funding acquisition.

## Data, code, and materials availability

The single-cell RNA sequencing data generated in this study have been deposited in the Gene Expression Omnibus (GEO) under accession number GSE317554. Code used to reproduce the single-cell RNA sequencing analyses and to generate the associated figures is publicly available at GitHub (https://github.com/AlmogAngel/GFPmilk). All other data are available in the manuscript or the supplementary materials. No new materials were generated that require material transfer agreements.

## Supplementary Materials

Supplementary Figs. 1-6 and Table S1

