## Supplementary Information for "Breastfeeding Enables Systemic Transfer of Functional Maternal Immune Cells to Neonates"

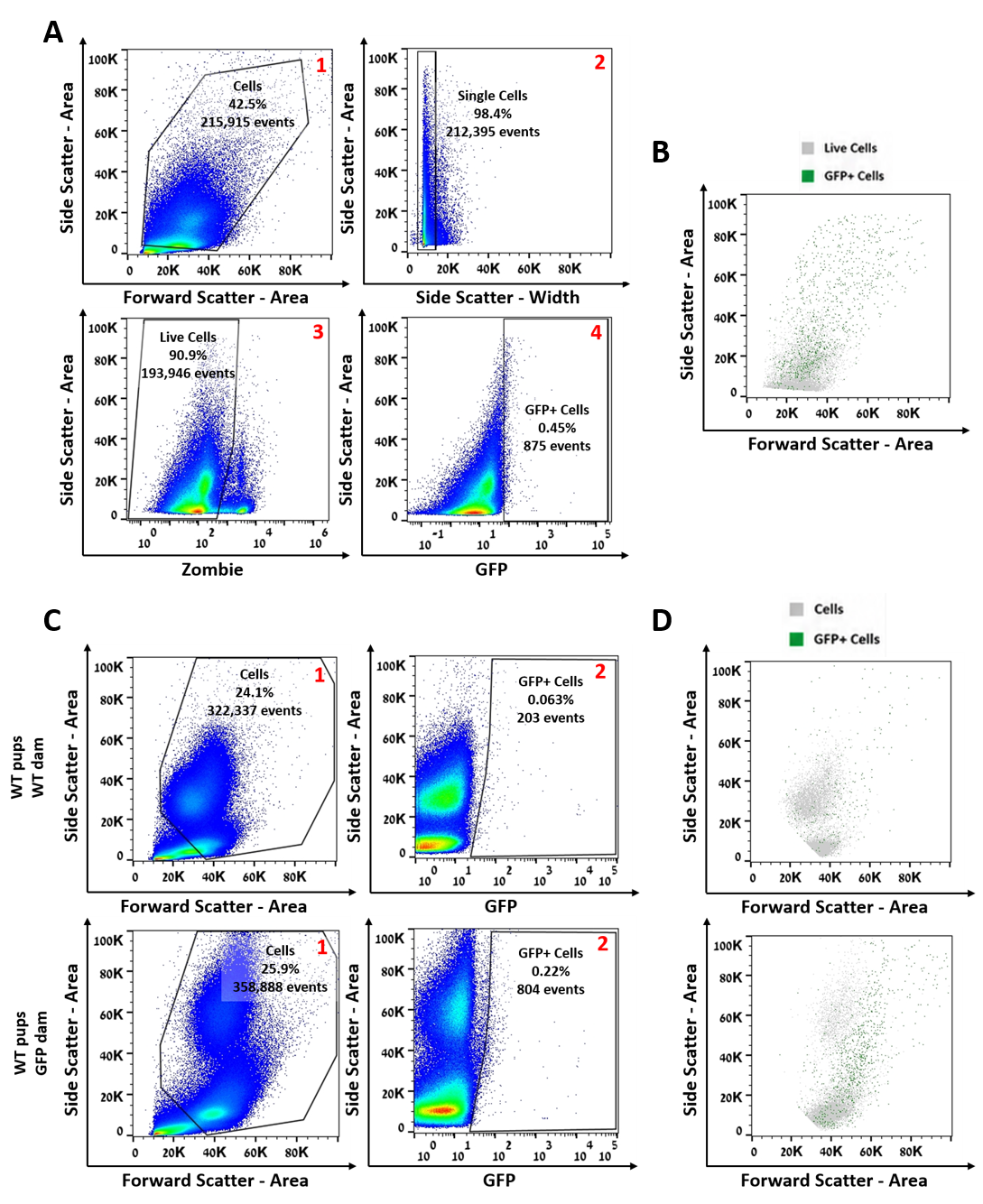

**Supplementary Figure 1. Flow cytometry gating strategy for identification of breastmilk-derived maternal GFP⁺ cells in neonatal blood and tissues. (A)** Representative gating strategy used to identify GFP⁺ maternal cells in blood and peripheral organs. Approximately 500,000 total events were acquired for each sample before sequential gating. All acquired events were first displayed on a forward scatter (FSC-A) versus side scatter (SSC-A) plot, and cellular events were selected while excluding residual debris (gate 1). Single cells were subsequently identified using side scatter area versus side scatter width (SSC-A versus SSC-W) to exclude doublets (gate 2). Viable cells were identified by exclusion of Zombie Violet-positive events (gate 3). GFP⁺ cells were then defined based on fluorescence intensity relative to unstained control (gate 4). The same gating strategy was applied to all tissues analyzed. Red blood cell (RBC) lysis was performed prior to flow cytometric analysis of blood, liver, and lung cell suspensions, whereas brain, spleen, and kidney cell suspensions were analyzed without RBC lysis. Percentages indicate the fraction of events retained at each gating step. **(B)** Representative back-gating of GFP⁺ events from the gating strategy shown in (A). Gray dots represent the live cell population (gate 3), whereas green dots represent GFP⁺ cells (gate 4), illustrating the distribution of GFP⁺ events within the live-cell population**. (C)** Representative flow cytometry plots from the sorting of peripheral blood of WT pups nursed by WT dams (top) or GFP⁺ dams (bottom). Approximately 1.5 million total events were acquired for each sample. GFP⁺ events are detected in WT pups nursed by GFP⁺ dams but are rare in pups nursed by WT dams, consistent with the maternal origin of these circulating GFP⁺ cells. Percentages indicate the proportion of GFP⁺ cells among live, singlet leukocytes. **(D)** Representative back-gating of GFP⁺ events from the peripheral blood analysis shown in **(C)**. Gray dots represent the total cell population (gate 1), whereas green dots represent GFP⁺ cells (gate 2), demonstrating the localization of GFP⁺ events within the overall cellular population. Each flow cytometry experiment was performed on pooled peripheral blood collected from a donor litter comprising five to seven nursed pups.

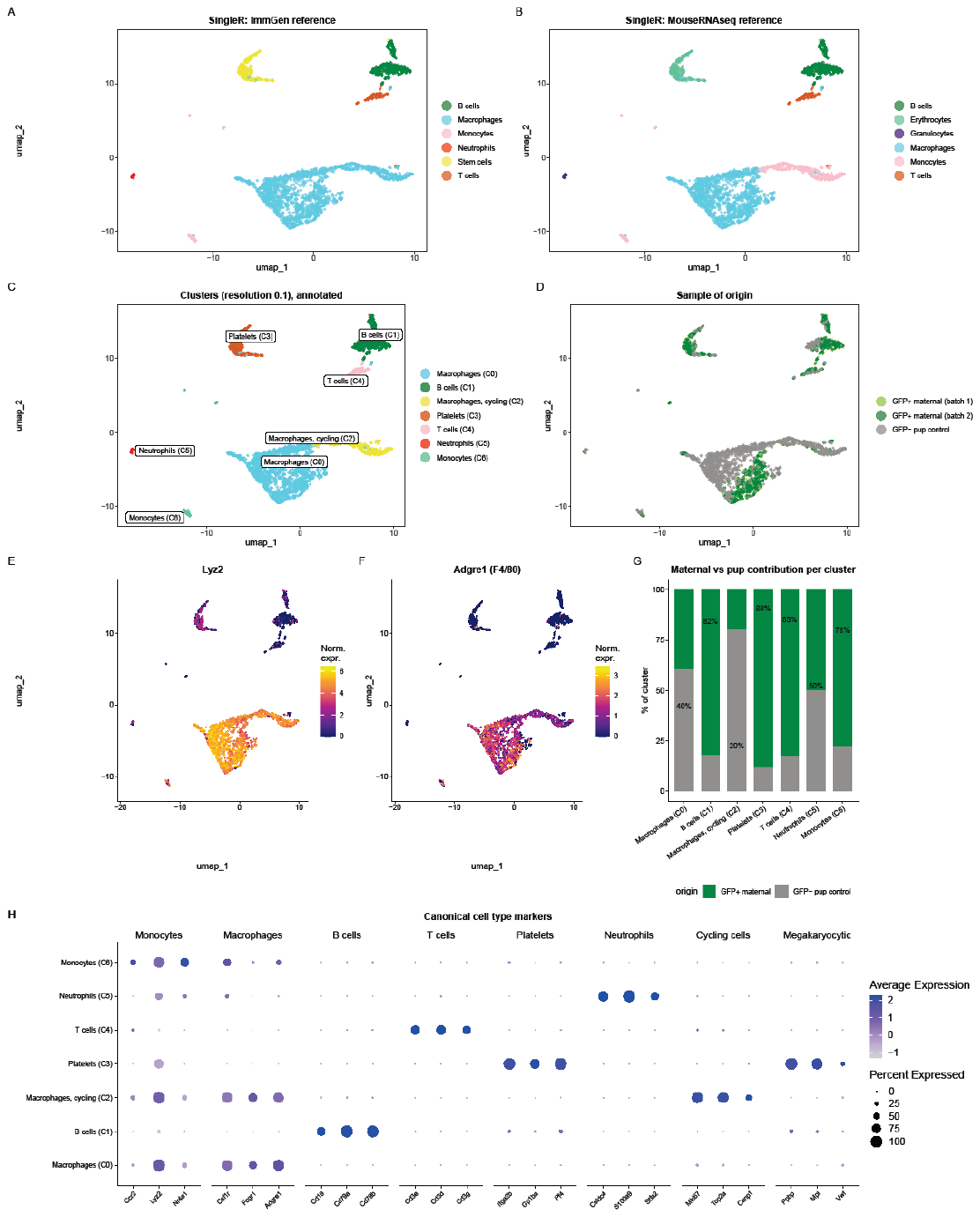

**Supplementary Figure 2.** **Cell type annotation of circulating blood cells.** (A, B) SingleR annotation of the clustered cells using the ImmGen (A) and MouseRNAseq (B) reference datasets. (C) UMAP of the seven clusters identified at resolution 0.1, labelled by assigned cell type. (D) The same UMAP colored by sample of origin, showing the contribution of each GFP⁺ maternal replicate and of the GFP⁻ pup control. (E, F) Normalized expression of the macrophage markers Lyz2 (E) and Adgre1/F4/80 (F), shown on a color scale from low (purple) to high (yellow). (G) Proportion of each cluster derived from GFP⁺ maternal samples versus GFP⁻ pup controls; the value above each bar is the maternal fraction. (H) Dot plot of canonical cell type markers across the seven clusters. Marker categories are monocytes (Ccr2, Lyz2, Nr4a1), macrophages (Csf1r, Fcgr1, Adgre1), B cells (Cd19, Cd79a, Cd79b), T cells (Cd3e, Cd3d, Cd3g), platelets (Itga2b, Gp1ba, Pf4), neutrophils (Cstdc4, S100a9, Stfa2), cycling cells (Mki67, Top2a, Cenpf) and megakaryocytic transcripts (Ppbp, Mpl, Vwf). Megakaryocytic transcripts are shown as a marker category and label the platelet cluster, as expected for platelets derived from megakaryocytes; no separate megakaryocyte cluster was identified. Dot size indicates the percentage of cells expressing each marker and color intensity the average expression level.

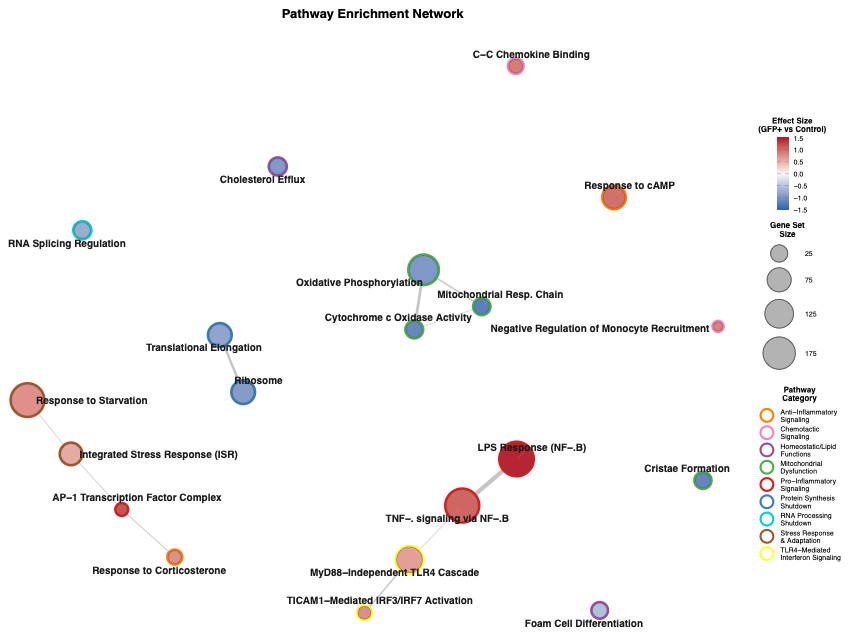

**Supplementary Figure 3. Gene enrichment network of maternal macrophage transcriptional programs.**

Network visualization of pathways significantly enriched or depleted in GFP⁺ maternal macrophages relative to GFP⁻ control macrophages. Nodes represent individual pathways, sized by gene-set size and colored by effect size (GFP⁺ vs control). Edges indicate gene overlap between pathways. Pathways cluster according to shared genes and functional themes, revealing discrete transcriptional modules-including inflammatory signaling, stress adaptation, metabolic/mitochondrial suppression, and RNA-processing shutdown-supported by largely non-overlapping gene sets.

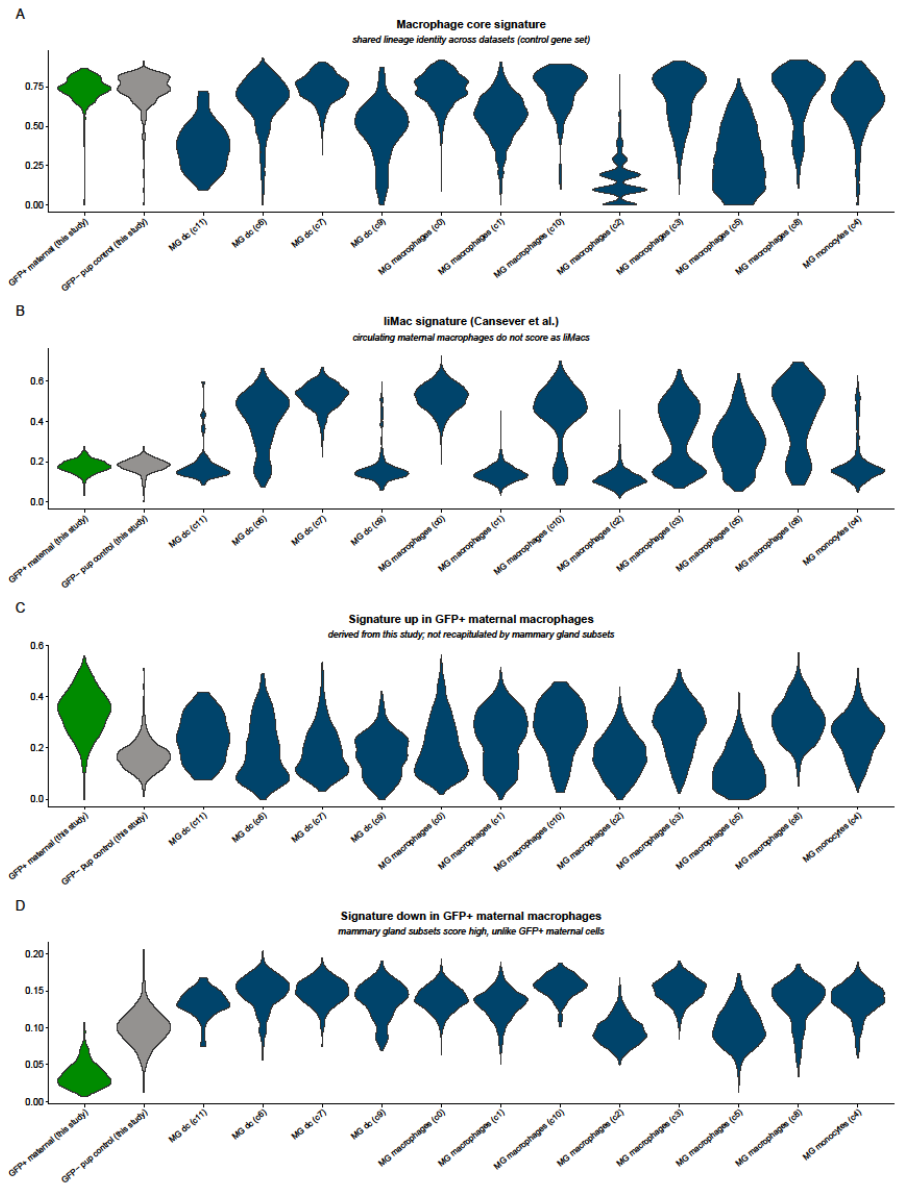

**Supplementary Figure 4. Circulating maternal macrophages are transcriptionally distinct from lactation-associated macrophages.** Macrophages profiled in this study (GFP⁺ maternal and GFP⁻ pup control) were compared with myeloid cells from the published lactating mammary gland dataset. Mammary gland populations are labelled by the cell identity assigned by SingleR, as macrophage, monocyte or dendritic cell subsets, with the corresponding cluster index in parentheses, rather than by cluster number alone. Each panel shows per-cell UCell scores for one gene set, plotted as violins across all groups. **(A)** A macrophage core gene set, scored comparably by circulating macrophages and by most mammary gland macrophage subsets, confirming shared lineage identity. **(B)** The liMac gene set defined in the original study; circulating maternal and control macrophages score low, whereas several mammary gland macrophage subsets score high. **(C)** The signature upregulated in GFP⁺ maternal macrophages relative to pup control macrophages in this dataset; GFP⁺ maternal cells are the only group in which it is enriched. **(D)** The signature downregulated in GFP⁺ maternal macrophages, for which GFP⁺ maternal cells score lowest of all groups. No mammary gland subset reproduces the paired pattern of high up-signature and low down-signature scores that distinguishes circulating maternal macrophages, supporting the conclusion that they represent a distinct macrophage state rather than a mobilized form of liMacs.

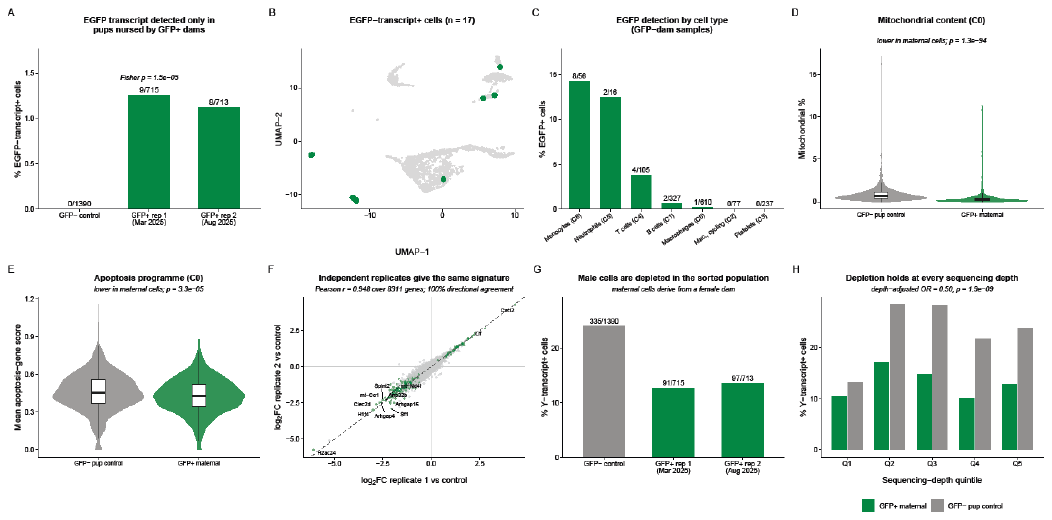

**Supplementary Figure 5. Transgene transcript detection, viability and replicate concordance in the single-cell dataset. (A)** Percentage of cells with detectable EGFP transcript in each sample; numbers above the bars give EGFP⁺ cells over total cells analysed. EGFP transcript was recovered from both GFP⁺ replicates and from none of the 1,390 control cells (Fisher exact test, p = 1.5×10⁻⁵). **(B)** UMAP showing the position of the 17 EGFP-transcript⁺ cells (green) among all profiled cells (grey). **(C)** Percentage of EGFP-transcript⁺ cells within each annotated cell type, restricted to pups nursed by GFP⁺ dams; numbers above the bars give EGFP⁺ cells over cells in that cluster. **(D)** Mitochondrial content and **(E)** mean apoptosis-gene score in GFP⁺ maternal versus GFP⁻ control macrophages (cluster C0); boxes show the median and interquartile range. Both are lower in maternal cells, arguing against the recovery of dying cells. **(F)** Concordance of the differential expression signature between the two independent GFP⁺ biological replicates, each compared against the control sample. Each point is a gene; green points are significant in both replicates and the dashed line is the identity. **(G)** Percentage of cells carrying transcripts from the Y-linked genes Ddx3y, Eif2s3y, Uty or Kdm5d in each sample; numbers above the bars give Y-positive cells over total cells. **(H)** The same percentage stratified by quintile of sequencing depth, showing that the depletion of male cells in the sorted population is not explained by the greater depth of the control libraries.

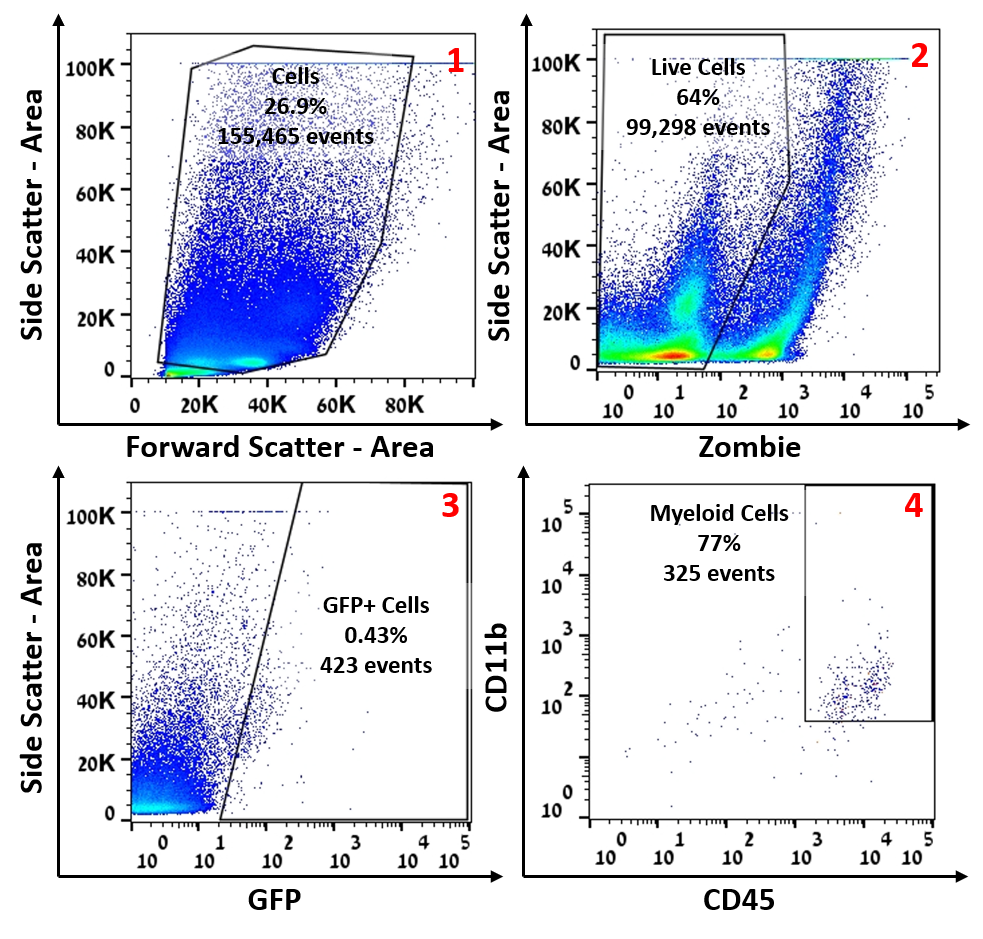

**Supplementary Figure 6. Flow cytometry gating strategy for sorting of GFP⁺ myeloid cells prior to qPCR analysis.** Representative sequential gating strategy applied to peripheral blood of WT pups nursed by GFP⁺ dams for isolation of maternal GFP⁺ cells by fluorescence-activated cell sorting (FACS) and subsequent quantitative PCR (qPCR) validation. The identical sequential gating strategy was applied to both experimental and control samples; however, for downstream qPCR, GFP⁺CD45⁺CD11b⁺ myeloid cells were sorted from pups nursed by GFP⁺ dams, whereas GFP⁺ cells were sorted from pups nursed by WT dams to obtain sufficient cell numbers. Gate 1: all acquired events were first displayed on a forward scatter area (FSC-A) versus side scatter area (SSC-A) plot; cellular events were selected while excluding debris. Gate 2: viable cells were identified by exclusion of Zombie Violet-positive events on SSC-A versus Zombie plot. Gate 3: GFP⁺ cells were identified based on fluorescence intensity on SSC-A versus GFP plot relative to an unstained control. Gate 4: among GFP⁺ events, myeloid cells were defined as CD45⁺CD11b⁺ on a CD11b versus CD45 plot. Percentages indicate the proportion of events retained at each sequential gating step. Red blood cell lysis was performed prior to analysis of all blood samples.

**Supplementary Table 1. Annotation of gene sets used for pathway enrichment analysis.**

| Pathway Name | Pathway ID | Dataset |
| --- | --- | --- |
| TNFa signaling via NF-kB | HALLMARK_TNFa_SIGNALING_VIA_NFKB | MSigDB Hallmark |
| AP-1 Transcription Factor Complex | GOCC_TRANSCRIPTION_FACTOR_AP_1_COMPLEX | Gene Ontology - Cellular Component |
| LPS Response (NF-kB) | GSE14769_UNSTIM_VS_40MIN_LPS_BMDM_DN | ImmuneSigDB (GEO dataset) |
| Response to cAMP | GOBP_RESPONSE_TO_CAMP | Gene Ontology - Biological Process |
| Response to Corticosterone | GOBP_RESPONSE_TO_CORTICOSTERONE | Gene Ontology - Biological Process |
| MyD88-Independent TLR4 Cascade | REACTOME_MYD88_INDEPENDENT_TLR4_CASCADE | Reactome |
| TICAM1-Mediated IRF3/IRF7 Activation | REACTOME_TICAM1_DEPENDENT_ACTIVATION_OF_IRF3_IRF7 | Reactome |
| Integrated Stress Response (ISR) | GOBP_INTEGRATED_STRESS_RESPONSE_SIGNALING | Gene Ontology - Biological Process |
| Response to Starvation | GOBP_RESPONSE_TO_STARVATION | Gene Ontology - Biological Process |
| C-C Chemokine Binding | GOMF_C_C_CHEMOKINE_BINDING | Gene Ontology - Molecular Function |
| Negative Regulation of Monocyte Recruitment | GOBP_NEGATIVE_REGULATION_OF_MONOCYTE_CHEMOTAXIS | Gene Ontology - Biological Process |
| Mitochondrial Resp. Chain | HP_MITOCHONDRIAL_RESPIRATORY_CHAIN_DEFECTS | Human Phenotype Ontology |
| Cytochrome c Oxidase Activity | GOBP_MITOCHONDRIAL_ELECTRON_TRANSPORT_CYTOCHROME_C_TO_OXYGEN | Gene Ontology - Biological Process |
| Cristae Formation | REACTOME_CRISTAE_FORMATION | Reactome |
| Oxidative Phosphorylation | GOBP_OXIDATIVE_PHOSPHORYLATION | Gene Ontology - Biological Process |
| Translational Elongation | GOBP_TRANSLATIONAL_ELONGATION | Gene Ontology - Biological Process |
| Ribosome | GOCC_RIBOSOME | Gene Ontology - Cellular Component |
| Cholesterol Efflux | GOBP_POSITIVE_REGULATION_OF_CHOLESTEROL_EFFLUX | Gene Ontology - Biological Process |
| Foam Cell Differentiation | HP_FOAM_CELLS | Human Phenotype Ontology |
| RNA Splicing Regulation | GOBP_NEGATIVE_REGULATION_OF_RNA_SPLICING | Gene Ontology - Biological Process |
